# Single cell RNA-seq in regenerative and fibrotic biomaterial environments defines new macrophage subsets

**DOI:** 10.1101/642389

**Authors:** Sven D. Sommerfeld, Christopher Cherry, Remi M. Schwab, Liam Chung, David R Maestas, Philippe Laffont, Julie E. Stein, Ada Tam, Franck Housseau, Janice M. Taube, Drew M. Pardoll, Patrick Cahan, Jennifer H. Elisseeff

## Abstract

Macrophages play diverse roles in the immune response to infection, cancer, and wound healing where they respond to local environmental signals, yet identification and phenotypic characterization of functional subsets *in vivo* remains limited. We performed single cell RNA sequencing analysis on differentiated macrophages sorted from a biologic matrix-induced regenerative environment versus a synthetic biomaterial foreign body response (FBR), characterized by T_H_2/interleukin (IL)-4 and T_H_17/IL-17, respectively. In the regenerative environment, unbiased clustering and pseudotime analysis revealed distinct macrophage subsets responsible for antigen presentation, chemoattraction, and phagocytosis, as well as a small population with expression profiles of both dendritic cells and skeletal muscle. In the FBR environment, we identified a CD9^hi+^IL-36γ^+^ macrophage subset that expressed T_H_17-associated molecules characteristic of certain auto-immune responses that were virtually absent in mice lacking the IL-17 receptor. Surface marker combinations including CD9 and CD301b defined macrophage fibrotic and regenerative subsets enabling functional assessment and identification in human tissue. Application of the terminal macrophage subsets to train the SingleCellNet algorithm and comparison to human and mouse macrophages in tumor, lung, and liver suggest broad relevance of macrophage classification. These distinct macrophage subsets demonstrate previously unrecognized myeloid phenotypes involved in different tissue responses and provide new targets for potential therapeutic modulation of certain pathologic states and tissue repair.

## Introduction

Macrophages are immune cells of myeloid lineage that maintain tissue homeostasis and participate in host defense. They remove cell debris, recycle, and clear apoptotic cells during tissue homeostasis and remodeling. As members of the innate immune system, macrophages sense and respond to infection, cancer, and tissue damage by cytokine and growth factor secretion and phagocytic activity. They are required for tissue regeneration and repair in addition to host defense. Through their cytokine profile and antigen presentation capacity, macrophages can engage and influence the adaptive immune system. For example, tumor-associated macrophages are a component of the tumor immune microenvironment where depending on their phenotype, they promote an immunosuppressive program that supports tumor growth or alternatively prime anti-tumor T cells to promote tumor regression (*1, 2*). Similarly, the wound associated macrophage phenotype can determine repair and fibrosis outcomes depending on T cell effector state (*3-5*).

Macrophages execute varied programs through their functional diversity and plasticity. They are highly specialized depending on tissue environment; i.e., alveolar macrophages in the lung (*6*), Kupffer cells in the liver (*7*), and microglial cells in the brain (*8*). Perturbations to the environment, particularly associated with certain disease states, alter their phenotype and function from normal tissue imprinting. The local tissue environment can influence macrophage phenotype beyond their developmental lineage. For example, environmental factors regulate macrophage epigenetics to control progenitor differentiation and reprogramming of already differentiated macrophages (*9*). In the tissue response to trauma and foreign bodies, macrophages play successive roles from initial inflammation to innate defense and resolution, resulting in tissue repair or chronic inflammation and fibrosis depending on environmental signals (*10*).

Macrophage functional heterogeneity is key to their ability to respond to diverse environments and cues. Yet this complexity in activity and gene expression is not reflected by current phenotyping and nomenclature dogma. Early attempts to classify macrophages resulted in the M1/M2 nomenclature to define the pro-inflammatory, interferon gamma (IFNγ) activated macrophage (M1) versus the “alternatively IL(Interleukin)-4 activated” macrophage (M2) (*11, 12*). Following this, a spectrum model of macrophage activation was proposed (*13*). In recent years, a common framework for macrophage activation has been suggested that considers a set of surface and genetic markers including CD206 as an “M2 marker” and CD86 as an “M1 marker” (*14*). Recent technological advances, however, including single cell RNA sequencing (scRNAseq) and mass cytometry suggest that archetypal *in vitro* phenotypes rarely overlap with those found in physiological conditions (*15, 16*). These approaches open the door to the refined characterization of macrophage populations that reflects the complexity of their phenotype and function in normal and pathological environments.

Biomaterials generate tissue microenvironments that can reproducibly induce specific macrophage phenotypes. Biological scaffolds, derived from the extracellular matrix (ECM) of tissues, promote a pro-regenerative tissue microenvironment that induces tissue repair through increased expression of IL-4. The repair capacity of these materials correlates with a T_H_2 T cell response that directs polarization to a traditionally-defined M2 macrophage in combination with a reduction of CD86 expression (*3, 4*). Synthetic materials induce a foreign body response (FBR) that has been associated with inflammatory M1-type macrophages and development of a fibrotic capsule (*17, 18*). While the M1-type inflammatory macrophage has been conventionally associated with a T_H_1 response, we have found that the FBR and the associated macrophage phenotype occurs in a type 17 immune environment that includes IL-17 production by innate lymphocytes, γδ T cells, and T_H_17 T cells. IL-17 signaling is required for the fibrosis associated with the FBR, though the level of IL-17 expression may vary depending on the chemical and physical properties of the materials (*19*). Synthetic and biological materials, therefore, serve as a model for Type 2 and Type 17 tissue immune microenvironments where the associated macrophage phenotype can be studied.

Here, we used scRNAseq to characterize macrophages isolated from a murine model of tissue repair versus fibrosis using model biomaterial tissue environments. Unbiased clustering algorithms revealed diverse populations of macrophages. Subsequent computational analysis identified phenotypic properties of the macrophage clusters. We identified inflammatory macrophages unique to regenerative and fibrotic environments with muted inflammatory and highly inflammatory signatures. We also defined phenotypically and functionally a phagocytic macrophage population exclusive to the reparative environment. Finally, we identified a unique CD9^hi+^IL-36γ^+^ fibrotic macrophage population in the FBR response with signatures of type 17 immune responses and autoimmunity. We also identified surface markers that defined the novel macrophage clusters and validated their ability to separate macrophage subsets experimentally by flow cytometry and immunofluorescence. Finally, we validated the broader relevance of the macrophage subsets in human pathologies including histiocytosis and fibrosis.

## Results

### Single cell profiling of F4/80^hi+^ macrophages in regenerative versus fibrotic tissue microenvironments

To model tissue microenvironments with activated and heterogeneous macrophage populations, we selected biomaterials that are used clinically and induce divergent immune and tissue responses. Biological scaffolds derived from the extracellular matrix of tissues promote healing of muscle (*20*) and induce a Type 2 (M2/T_H_2) immune microenvironment (*4*). We selected a urinary bladder matrix (UBM), with properties similar to other ECM-derived materials, that is used clinically in wound healing (*21*) and hernia repair (*22*). Polycaprolactone (PCL) was selected as a model synthetic biomaterial that stimulates a fibrotic response (*23*) and a Type 17 (M1/T_H_17) immune microenvironment (*19*). When characterizing macrophages in these biomaterial tissue microenvironments, CD86 (a ligand for CD28 and CTLA4) and CD206 (a scavenger receptor) expression is analyzed on differentiated macrophages (CD45^+^CD64^+^F4/80^hi+^) (*24*) to characterize polarization. We sorted this population of differentiated macrophages for single cell analysis one week after biomaterial implantation and applied them to the 10X single cell RNA sequencing platform (Fig. 1A). This resulted in the capture of ∼7,200 macrophages with an average read depth of ∼50,000 reads per cell across ∼13,000 genes with over 4,000 median unique molecular identifiers (UMI) counts per cell (Supplementary Fig. 1 and Supplementary Table 1). By condition, the total number of macrophages captured was 3,343 from UBM, 2,919 from PCL, and 876 from saline.

**Figure 1.**
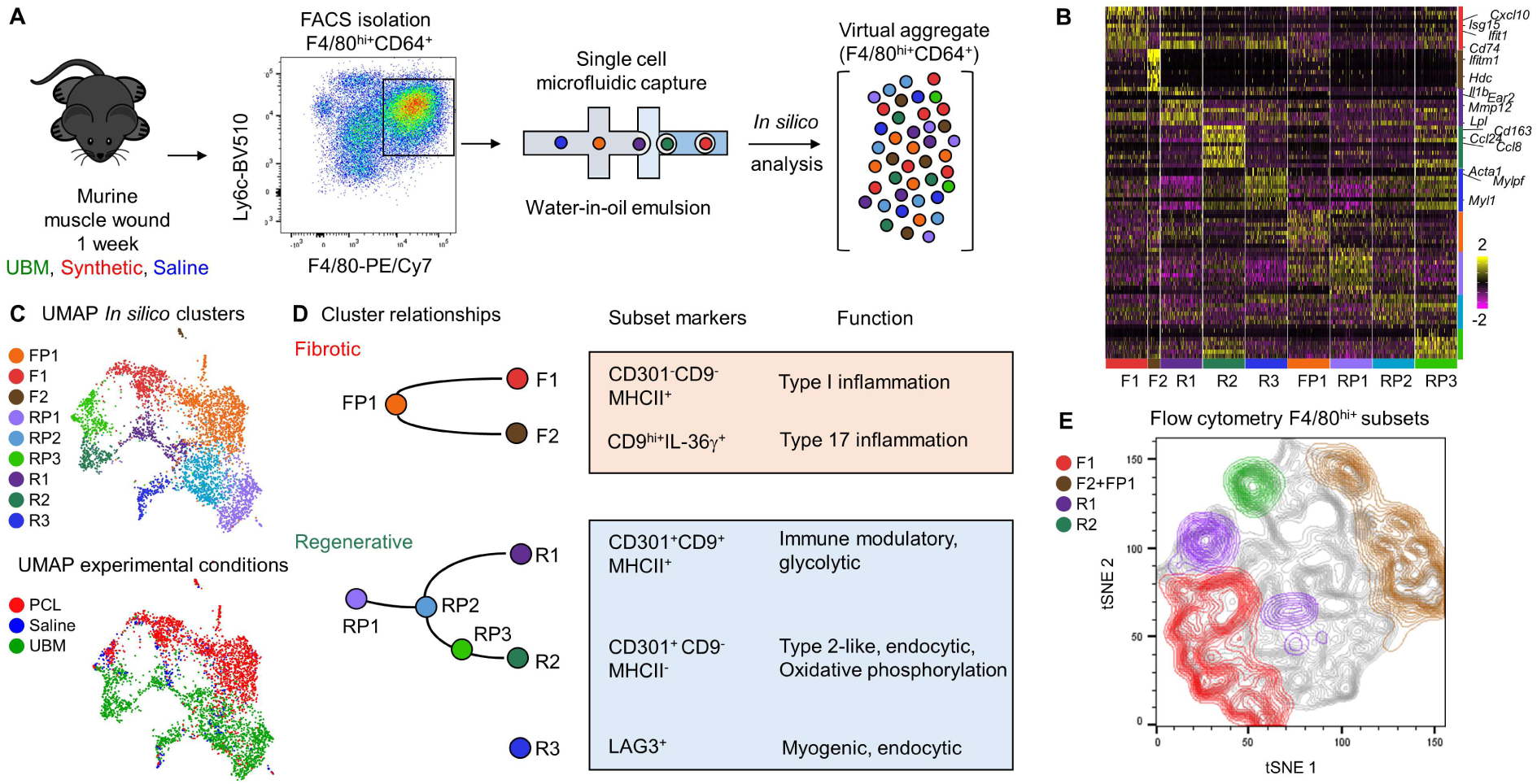
Single cell characterization of macrophages in fibrotic and regenerative microenvironments. (A) Experimental overview. A virtual aggregate of macrophages in fibrosis and regeneration generated from single cell RNA-seq after sorting of F4/80^hi+^CD64^+^ cells isolated from murine volumetric muscle injuries at 1 week, treatment with biomaterials UBM (regenerative), synthetic (fibrotic), or saline (control). (B) Heatmap of differentially expressed genes. Up to 200 cells per cluster are shown, ordered by cluster, with the top 10 differentially expressed genes. Functionally relevant genes from terminal clusters are annotated. (C) Dimensional reduction projection of cells onto two dimensions using uniform manifold projection approximation (UMAP). Cells are colored by experimental biomaterial condition (top) and computationally determined cluster (bottom). (D) Summary of cluster differentiation trajectories, markers, and biological functions generated by bioinformatics analysis. (E) A flow cytometry strategy informed by *in silico* determined markers including CD9, CD301b and MHCII differentiating *in vivo* macrophage subsets from UBM, synthetic. Subsets are colored equivalent to *in silico* clusters, back gated into tSNE projection.

### Unbiased clustering and pseudo-time trajectories reveal functional diversity that correlates with biomaterial tissue microenvironment

Unbiased clustering algorithms categorized macrophages into clusters based on global gene expression patterns. We first computationally pooled macrophages from regenerative (UBM), fibrotic (PCL), and control (saline) conditions to create a virtual aggregate. Cells with reduced signal after scaling were removed from the analysis (Supplementary Fig. 2), leaving 9 computationally determined clusters. The clusters were largely enriched for regenerative or fibrotic macrophages with differential expression analysis confirming distinct gene expression profiles (Fig. 1B). UMAP (uniform manifold approximation and projection), a dimensional reduction algorithm, grouped cells by cluster, indicating that UMAP and the clustering algorithm agreed on the similarity of cell phenotypes (Fig. 1C).

The experimental origin of each cell was then superimposed on the UMAP plot so that the enrichment of cells from regenerative or fibrotic microenvironment could be identified across all cell clusters (Fig. 1C). Cell clusters were distinguished by experimental conditions, so we termed macrophages from the UBM tissue microenvironment regenerative associated clusters (RACs) and those from the PCL tissue microenvironment fibrosis associated clusters (FACs). Potential batch effects were removed by scaling on each experimental condition. Groups of macrophages isolated from saline-treated wounds were found at the interfaces between fibrotic and regenerative macrophages. This intermediate profile is supported by flow cytometry data suggesting that macrophages in a muscle wound without a biomaterial have characteristics of fibrotic and regenerative microenvironment (Fig. 2A). Similar cell numbers were sequenced from the fibrotic and regenerative conditions, but two-thirds of the macrophage cell clusters were RACs suggesting there is increased complexity of macrophage phenotypes in the regenerative tissue microenvironment.

**Figure 2.**
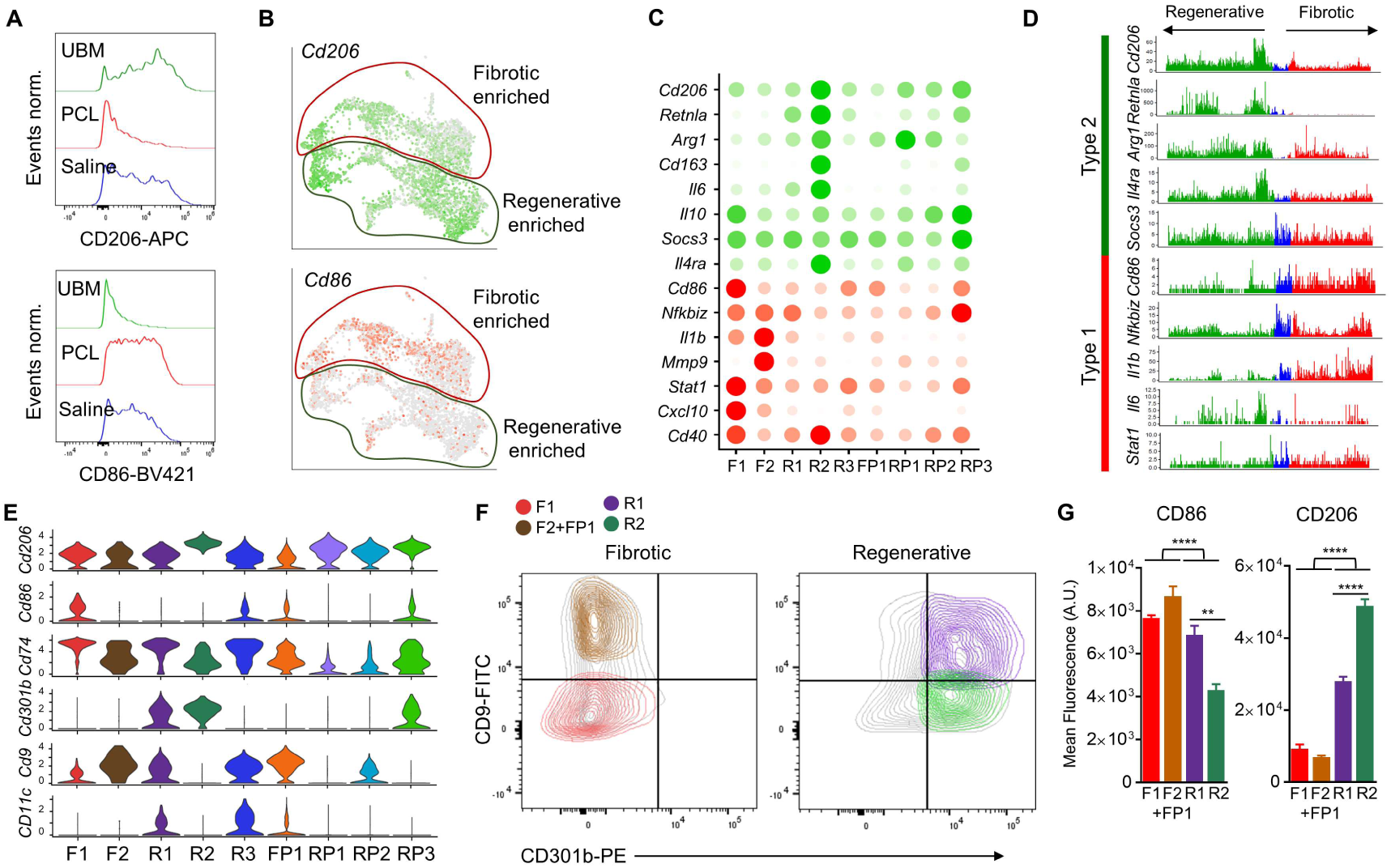
ScRNAseq reveals surface markers that discriminate diverse regenerative and fibrotic macrophage clusters. (A) Histogram of CD86 and CD206 expression of bulk macrophages from regenerative, fibrotic, and saline microenvironments (green, red, blue) by flow cytometry. (B) Feature plots of *in silico Cd86* (bottom) and *Cd206* (*Mrc1*, top) expression superimposed on UMAP plots of cells from scRNAseq. Circled borders mark regions enriched for cells from the regenerative or fibrotic experimental condition. (C) Cluster expression of canonical M1 (red) and M2 (green) markers. Expression levels are given as cluster averages normalized to the maximum value per gene. (D) Gene expression (UMI count) of M1 and M2 markers per single cell. Cells are ordered and colored by condition (regenerative as green, saline as blue, and fibrotic as red). (E) Violin plots of cluster *in silico* gene expression for surface markers. Surface markers were identified by differential expression analysis. (F) Flow cytometry gating strategy using CD9 and CD301b differentiates fibrotic and regenerative macrophage subsets *in vivo*. (G) Mean fluorescence values indicate subset specific expression of activation markers CD86 and CD206 (n = 4 biologically independent, **p<0.005, ****p<0.0001).

To identify relationships between cell clusters and differentiation trajectories, we performed Slingshot pseudotime and RNA velocity analysis on the RACs and FACs. Precursor clusters (RP1, RP2, FP1) were selected based on similarities in gene expression in clusters across experimental conditions (Supplementary Fig. 3). While RNA velocity, which predicts cell movement on a ∼32 h timescale, confirmed movement of cells from RP2 towards R1 and supported the defined clusters. Pseudotime results indicate a branching lineage in both the RACs (R1, R2) and FACs (F1, F2) (Fig. 1D) with two functionally specialized terminal clusters in each condition. R3 was excluded from the pseudotime analysis due to its unique gene expression profile that included muscle-related genes.

To enable identification of the terminal regenerative and fibrotic macrophages, we determined surface marker combinations *in silico* that could differentiate subsets experimentally (Fig. 1D). We performed flow cytometry on cells isolated from the UBM, PCL, and saline treatment conditions using the computationally-identified cluster surface markers. The CD45^+^Ly6c^-^F4/80^hi^ cell populations from all conditions were concatenated together to create a t-Distributed Stochastic Neighbor Embedding (tSNE) plot containing a complex mixture of all macrophages. We then identified macrophages expressing the surface markers CD9 (a protein involved in cell adhesion, fusion, and motility), CD301b (a galactose C-type lectin), and MHCII in the aggregated data set to represent the *in silico* macrophage clusters. The four terminal clusters F1, F2 (and FP1), R1, and R2 could be separated in the aggregate, suggesting that the new populations can be readily identified experimentally using flow cytometry (Fig. 1E).

### Expression of canonical polarization markers CD86 and CD206 is distributed across macrophage clusters

We first explored the correlation of the unbiased single cell clusters with canonical M1 and M2 polarization markers. Flow cytometry analysis of macrophages confirmed the enrichment of CD206 in the regenerative condition and CD86 in the synthetic condition with saline or untreated wound exhibiting intermediate levels of both markers (Fig. 2A). Histograms were consistent with previous studies that found UBM treatment downregulates CD86 while CD206 remains constant and PCL slightly decreases CD86 and significantly decreases CD206 compared to saline treatment.

While scRNAseq supported the enrichment of *Cd206* across regenerative macrophages and *Cd86* across fibrotic macrophages, it could not discriminate between phenotypically distinct macrophages subsets. Expression levels of these two surface markers superimposed on the UMAP plot show neither *Cd86* nor *Cd206* correlated with the computationally determined clusters (Fig. 2B). We then compared the expression patterns of canonical polarization genes and *Cd206* and *Cd86* across the unbiased clusters of RACs and FACs and found similar disparities (Fig. 2C). In particular, cluster F1 had elevated expression of *Cd86*, but relatively low levels of other M1 genes *Il1b, Mmp9*, and *Nfkbiz*. Meanwhile, *Il1b* and *Mmp9* were expressed predominantly in fibrotic cluster F2 which had the lowest expression levels of *Cd86*. The strongest expression of M2 markers was found together with the highest expression of *Cd206* in R2, but R2 also had high expression of the M1 marker *Cd40*. Other clusters had elevated expression of the type 2 associated genes *Arg1* and *Socs3* despite reduced expression of *Cd206*.

Comparison of macrophage polarization markers on a per cell basis in the different experimental conditions also revealed a significant heterogeneity (Fig. 2D). *Retnla* was the only type 2 gene not expressed in the fibrotic macrophages. Expression of other type 2 genes, *Arg1* and *Socs3*, did not parallel *Cd206* expression on a cell-by-cell basis. At the same time, high levels of *Retnla, Arg1*, and *Socs3* expression were found in cells that did not express *Cd206*. There was a similar pattern of disparity in expression of *Cd86* and type 1 genes. *Cd86* expression did not correlate with *Nfkbiz*, and *Il1b* on a cell-by-cell basis. Many cells expressed high levels of *Cd86* expression in parallel with low levels of *Nfkbiz* and *Il1b*.

### CD9, CD301b, and MHCII expression identifies fibrotic and regenerative macrophage subsets

Since the single cell analysis confirmed that *Cd86* and *Cd206* expression did not differentiate phenotypic subsets, we explored alternative surface markers in the scRNAseq dataset. *In silico* assessment of surface markers revealed that differential expression of *Cd301b, Cd9*, and *Cd74* (encoding the MHCII-associated invariant chain) was sufficient to identify each of the macrophage clusters (Fig. 2E). We then tested these surface markers on bulk cell isolates from the tissue environments to confirm that the gene expression correlated with surface protein expression and could separate the macrophage populations using multiparametric flow cytometry. The proposed surface markers were able to discriminate the macrophage subsets corresponding to *in silico* determined clusters in the regenerative and fibrotic conditions (Fig. 2F) using the gating schematics in Supplementary Fig. 4, 5. The surface marker CD301b allowed nearly complete separation of macrophages unique to the regenerative microenvironment while CD9 and CD74 further differentiated macrophages into the multiple *in silico* predicted subsets (Fig. 2G). The CD9 and CD301b surface marker paradigm differentiates macrophage groups not equivalent to the commonly used cytokine-induced *in vitro* macrophage phenotypes (IL-4, IFNγ^+^LPS, IL-10) that all express high levels of CD9 (Supplementary Fig. 6).

### Computational phenotyping reveals limited inflammatory and phagocytic macrophages in the regenerative environment

We used pseudotime analysis to elucidate relationships between the RACs and found a branching differentiation trajectory with two terminally differentiated clusters (R1, R2) stemming from three precursor clusters (RP1-3) (Fig. 3A, B). Precursor RP3, a direct precursor to only R2, shared a similar but reduced phenotype to R2 based on differential gene expression (Supplementary Fig. 7). While R2 is composed entirely of macrophages derived from the regenerative condition, RP3 had a relatively high composition of saline derived macrophages (Supplementary Fig. 8). The RACs expressed *Il4ra* with R2 expressing the highest levels, correlating with the UBM induction of IL-4 and the macrophage response to the tissue environment induced by the biological scaffold.

**Figure 3.**
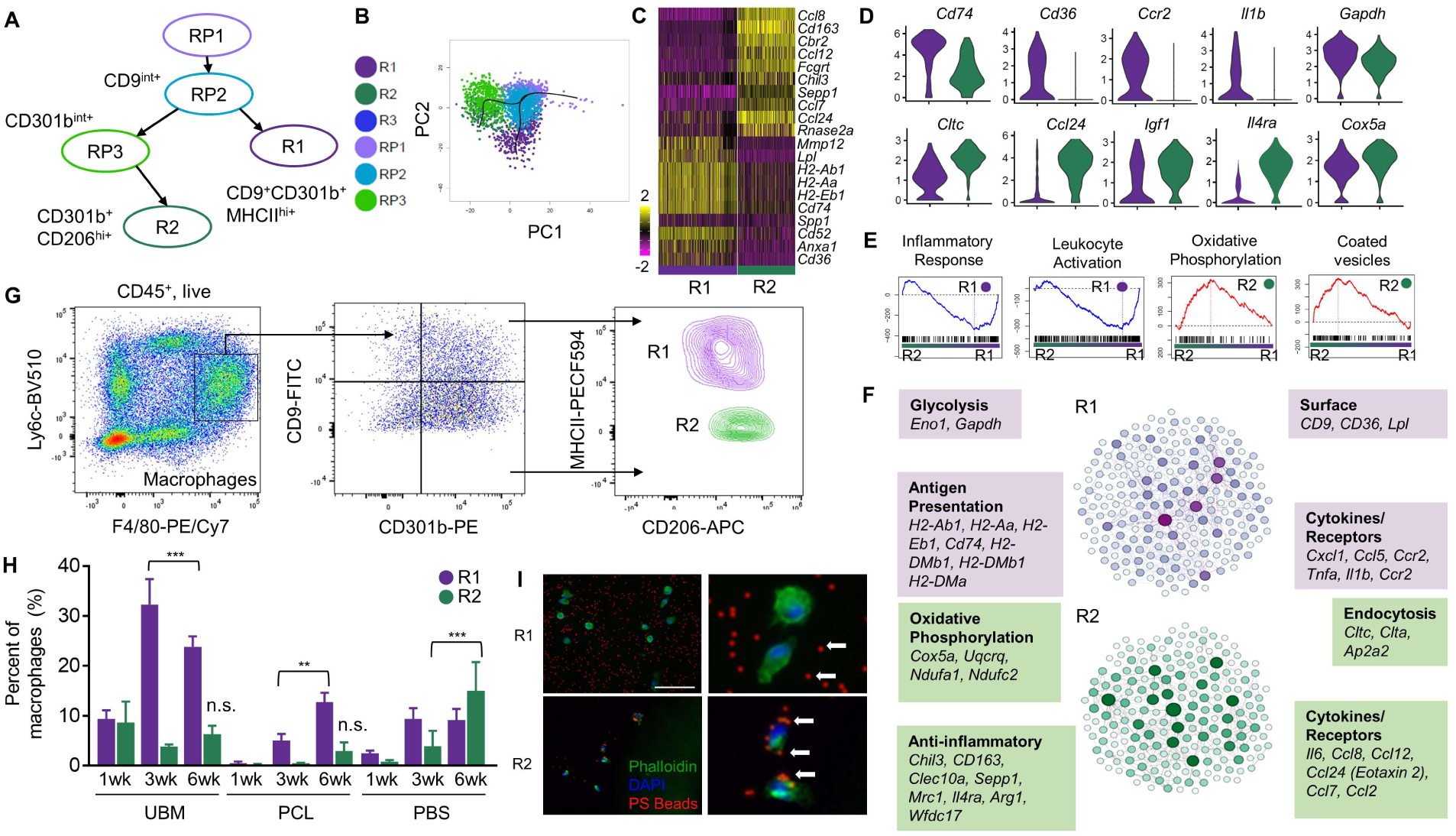
*In silico* RACs reveals distinct *in vivo* regenerative macrophage phenotypes including muted-inflammatory R1 and phagocytotic R2. (A) Predicted lineage schematic of RACs from Slingshot analysis. *In silico* predicted surface markers are shown. (B) Slingshot pseudotime trajectory of RACs shown on a principal component plot (PC1 vs. PC2). Cells are colored by cluster. (C) Heatmap of top 20 differentially expressed genes in a comparison of R1 and R2. (D) Violin plots for differentially expressed genes comparing R1 and R2. (E) Gene set enrichment comparing R1 and R2. Plots with higher peaks (red) indicates enrichment of gene sets in R2 while plots with negative peaks (blue) indicate enrichment of gene sets in R1. (F) Gene network plots of R1 and R2 generated. Nodes represent genes with connections generated by STRING metadata analysis. Sets of genes associated with specific functions are annotated. (G) Flow cytometry gating scheme validates *in vivo* protein marker combination for R1 and R2. Macrophages defined as F4/80^hi+^ from live, CD45^+^. (H) 1–6 weeks’ time course of R1 and R2 subsets in UBM, PCL, and saline microenvironments (n =4 biologically independent). Two-way analysis of variance with subsequent multiple testing p values are presented. (****p <0.0001). (I) Phagocytosis of flash red fluorescent microbeads by *ex vivo* cultured, sorted R1 and R2 macrophages, arrows indicate poly(styrene) bead locations (scale bar = 50 μm).

To characterize the phenotype and potential functional properties of R1 and R2, we compared outcomes of their differential expression, gene set enrichment analysis, and gene network analysis. Differential gene expression analysis of R1 showed upregulation of antigen-presenting capacity, inflammatory activity, and glycolysis, including *Cd74, Ccr2, Il1b*, and *Gapdh* (Fig. 3C, D). While R1 expression levels of the inflammatory genes *Il1b* and *Tnfa* were elevated when compared to other RACs, they were low when compared to the highly inflammatory FACs (Supplementary Fig. 9). Gene set enrichment of R1 also found elevation of inflammatory responses (Fig. 3E). R1 was enriched in leukocyte activation gene sets, suggesting that these macrophages play a role in communication and activation of the adaptive immune system. Finally, network analysis supported both the differential expression and gene set enrichment. R1 expressed gene modules associated with glycolysis (*Eno1, Gapdh*), antigen presentation (*H2* genes and *Cd74*), and inflammatory cytokines and chemokines (*Cxcl1, Ccr2, Ccl5, Tnfa*, and *Il1b)* (Fig. 3F).

In contrast to the inflammatory R1 macrophage subset, R2 expressed multiple genes associated with alternatively-activated or anti-inflammatory macrophages (Fig. 3C). Differential expression and network analysis both revealed enrichment for anti-inflammatory genes such as *Chil3, Cd163*, and *Mrc1* (gene encoding CD206) (Fig. 3D). In addition, R2 expressed high levels of *Ccl24* (*Eotaxin-2*), a chemokine for eosinophil attraction that is observed responding to ECM materials (*25*). Gene set enrichment and gene modules from network analysis also support a unique metabolic profile with expression of *Cox5a, Uqcrq, Ndufa1*, and *Ndufc2* (Fig. 3E, F). This profile supports R2 activation of oxidative phosphorylation compared to glycolysis in R1. R2 expression also included gene set enrichment and endocytic gene modules (*Cltc, Clta*, and *Ap2a2*) that suggest phagocytic activity in this subset.

### *In vivo* subtyping validates *in silico* predicted RAC phenotypes and microenvironment enrichment using CD9 and CD301b-based flow cytometry

*In silico* defined clusters R1 and R2 were validated *in vivo* by flow cytometry using a combination of CD301b, CD9 and MHCII on cells isolated from the UBM tissue microenvironment. While CD301b separated the terminal regenerative clusters R1 and R2 from progenitor clusters and R3, we found the R1 and R2 could be further defined as CD9^+^MHCII^+^ and CD9^-^MHCII^-^ respectively. A combination of both markers provided distinct separation of R1 from R2 for analysis and cell sorting (Fig. 3G). As predicted computationally, CD301b^+^CD9^+^ R1 and CD301b^+^CD9^-^ R2 macrophages express different levels of both MHCII and CD11c when quantified by surface marker expression using flow cytometry (Supplementary Fig. 10). R1 and R2 are distinguished by higher CD11c and lower CD206 expression respectively. These pronounced subset specific profiles elucidate earlier suggestions of complex phenotypes associated with the regenerative UBM microenvironment (*25, 26*).

To evaluate the kinetic evolution of the regenerative macrophage subsets in different experimental conditions, we performed flow cytometry on macrophages isolated from regenerative, fibrotic, and control VML microenvironments at 1, 3 and 6 weeks (Fig. 3H). Consistent with RNA velocity analysis (Supplementary Fig. 11), R1 increased significantly in the regenerative condition from 1 to 3 weeks while the R2 population maintained elevated levels with respect to the other conditions. As expected, the fibrotic condition had low expression of both R1 and R2 at all timepoints. The saline wound control showed an increase in both R1 and R2 at 3 weeks and R2 at 6 weeks, suggesting the UBM tissue microenvironment may be following an accelerated course of regeneration that is observed functionally with respect to an untreated wound.

Expression analysis predicted that R2 macrophages were phagocytic, an important function in wound clean up that is associated with tissue repair. Since the surface markers uniquely identified R2 by flow cytometry, we were able to sort pure populations of R1 and R2 and experimentally validate phagocytic activity. R1 and R2 macrophages sorted from a UBM-treated regenerative environment were evaluated for the ability to uptake fluorescent microbeads. The R2 (CD301b^+^CD9^-^) macrophage subset phagocytosed beads whereas R1 (CD301b^+^CD9^+^) macrophages did not internalize any beads (Fig. 3l). This result confirms the functional properties predicted by the gene expression analysis and provides surface markers that can specifically identify phagocytic macrophages.

### Genes associated with the local tissue microenvironment are found in macrophage cluster R3

The R3 RAC expressed the most unique gene signature compared to all other subsets with UBM treatment. Many of the top differentially expressed genes in this macrophage population were related to muscle tissue *Mylpf, Myl1, Acta1, Tnnc2*, and *Tnnt3* (Fig. 4A) and dendritic cells (*Lag3, Cd11c)* (*27*), Gene set enrichment and network analysis indeed supported this finding, with gene modules associated with skeletal muscle (*Myl4, Des, Ttn, Tnnc2, Tpm1*, and *Acta1*) (Fig. 4B) and strong enrichment of gene sets related to myogenesis and muscle function (Fig. 4C). Deeper analysis found that R3 cells also expressed genes characteristic of endocytosis and lysosome activity including moderate elevation of *Psap, Ctss, Hexb, Cd63*, and *Ctsz* (Fig. 4D). This is further supported by gene set enrichment finding enrichment of sets related to lysosomal and endocytic function and network analysis showing a module of genes associated with antigen presentation (*B2m, Cd74*, and *H2* genes). Two possible explanations emerge from these results. R3 muscle signature may be due to macrophages differentiating towards a muscle lineage and participating in building new muscle tissue since this is the tissue environment in which they are located. Alternatively, the single cell analysis is detecting endosomal mRNA in macrophages that have phagocytosed muscle cells during cell and tissue debris removal in the wound healing process.

**Figure 4.**
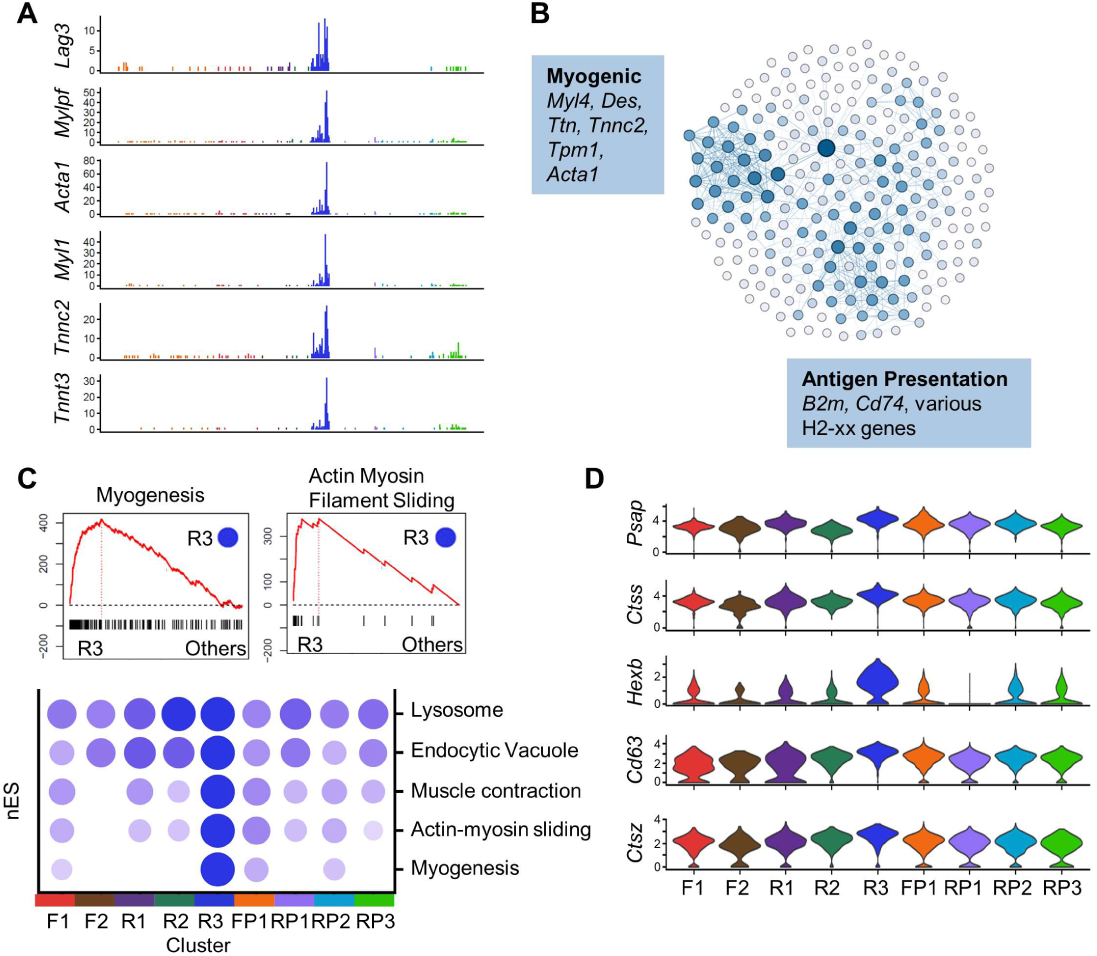
Cluster R3 expresses tissue specific genes. (A) UMI counts for genes overexpressed in R3. Genes shown here are associated with skeletal muscle function. (B) Violin plots of genes associated with endocytosis and lysosome activity. (C) Network analysis of F2 genes. Nodes are genes with edges connecting nodes representing connections in databases or literature. Modules of genes with associated functions are annotated. (D) Gene set enrichment on differential expression comparison of R3 to all other macrophages. Running enrichment plots (top) show high peaks when gene sets are overrepresented in R3 differentially expressed genes. The heatmap shows enrichment scores normalized across clusters for gene sets found enriched in R3.

### Computational phenotyping indicates FACs associated with inflammation and autoimmunity

We also applied pseudotime and RNA velocity analysis to determine FAC differentiation and polarization relationships (Supplementary Fig. 11). From pseudotime analysis, one precursor cluster (FP1) led to a branching trajectory with two terminal fibrosis-associated macrophage clusters, F1 and F2 (Fig. 5A). RNA velocity confirmed pseudotime results and continued polarization of F2 from the bulk macrophages. The F1 cluster expressed traditional markers of inflammation and genes associated with the interferon response including *Irf7, Il18*, and *Tlr2* (Fig. 5B). Gene sets for both IFNα and IFNγ response were enriched in F1 (Fig. 5C). Gene networks also showed modules associated with interferon response (*Stat1, Myd88, Irf7*, and *Tlr2*) and cytokines associated with inflammatory function (*Il18, Ccl4, Ccl7*, and *Cxcl10*) (Fig. 5D). While both macrophage populations have elements of inflammation, F1 and F2 exhibit significantly different expression profiles.

**Figure 5.**
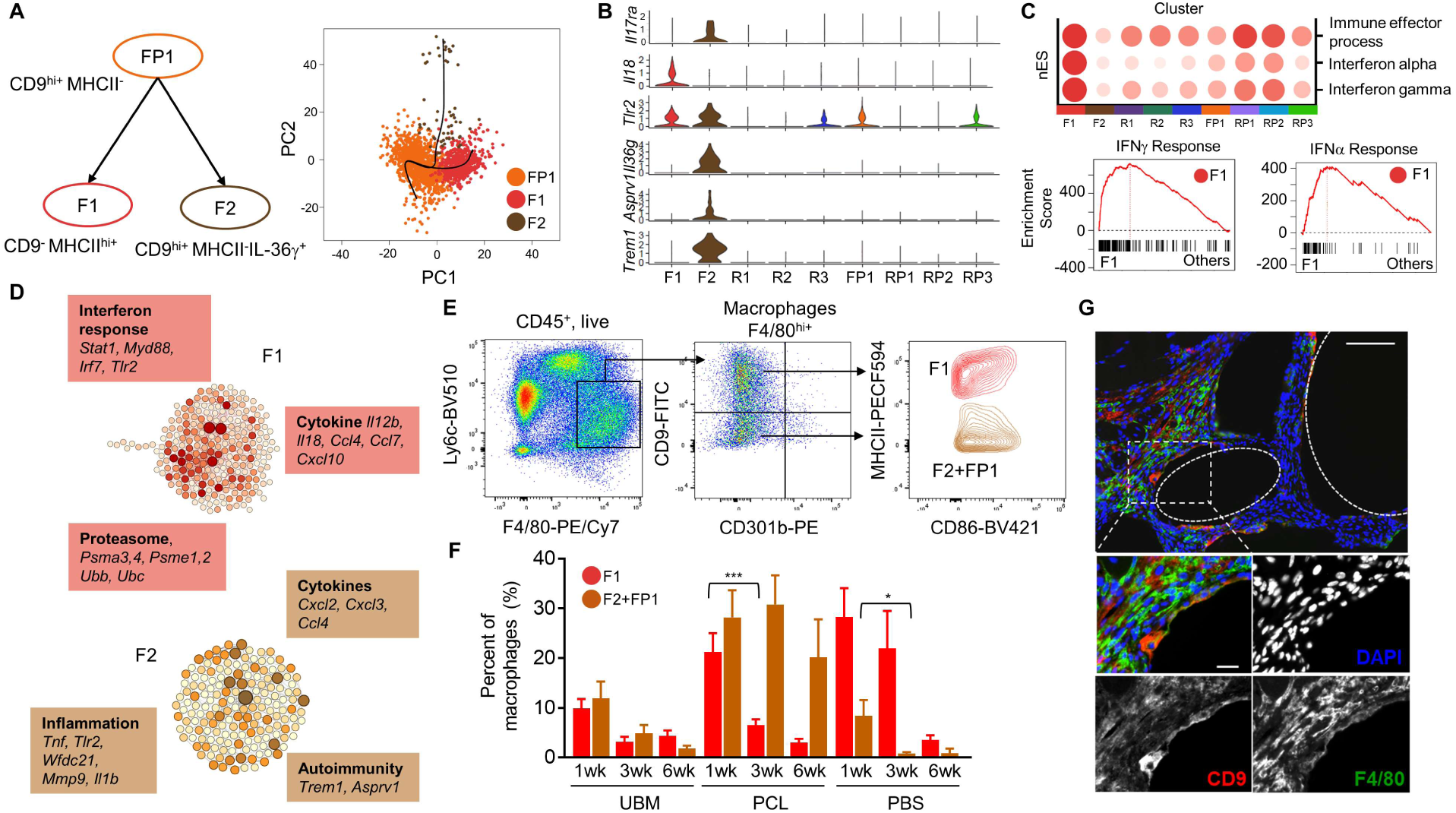
Fibrotic associated macrophages include distinct subsets F1 (MHCII^hi+^) and F2 (CD9^hi+^IL-36γ^+^). (A) Lineage schematic of RACs from Slingshot pseudotime analysis and trajectory including descriptive marker combination. Pseudotime trajectory is shown in a principal component plot (PC1 vs PC2). (B) Fibrotic subsets are distinguished by specific marker profile *in silico.* (C) Heatmap of gene set enrichment scores normalized across clusters for gene sets found upregulated in F1 and running gene set enrichment plots for the IFNγ and IFNα responses. (D) Gene network representation for relationships of differentially expressed genes in F1 (top) and F2 (bottom) by STRING metadata scores. (E) Flow cytometry gating strategy specific to F1 and (F2+FP1) from F4/80^hi+^ macrophages using CD9, MHCII, CD11c (F) Time course of F1 and (F2+FP1) subsets in UBM, PCL and saline microenvironments (n = 4 biologically independent). Two-way analysis of variance *p* values are presented (\**p* <0.05, \*\*\*\**p* <0.0001). (G) Immunofluorescence histology for CD9 (red) and F4/80 (green) at 1 week VML with synthetic material (scale bars = 100 μm and 25 μm, respectively).

While F2 was small, it had a distinct gene expression signature that included recently discovered cytokines and genes with limited characterization. While F2 expressed inflammatory markers *Slpi, Hdc, Tlr2*, and *Il1b* (Supplementary Fig. 12), it also expressed genes associated with autoimmunity *Il36γ, Trem1, Asprv1, and Il17ra* (Fig. 5B). The unique nature of this subset and possible functional relevance is exemplified in the expression of *Il36γ* (also known as *Il1f9*) that is found in lesions of skin psoriasis and participates in a positive feedback loop in type 17 immune responses (*28*). IL-17, the primary cytokine of Type 17 responses is also associated with fibrotic diseases not yet associated with autoimmunity including idiopathic pulmonary fibrosis (IPF) (*29, 30*), cardiac fibrosis (*31*), and the foreign body response suggesting a Type 17-autoimmune connection. The F2 cluster expressed increased *Il17ra*, further supporting the role of IL-17 in this subset and the macrophage responsiveness in the PCL tissue microenvironment.

To validate the experimentally the FAC clusters, we performed flow cytometry using the computationally defined surface marker strategy including CD9, CD301b, and MHCII (Fig. 5E and F). According to the computational analysis, the surface markers differentiate the F1 cluster but the F2 and FP1 clusters are not separated. Both F1 and F2+FP1 were elevated in the PCL fibrotic condition at 1 week with a significant reduction in F1 from 1 to 3 weeks. However, levels of F2+FP1 were consistently elevated, suggesting that F2 may be important in the development of pathological fibrosis. The UBM regenerative condition had low levels of both F1 and F2 at 1 week which was reduced at 3 and 6 weeks. The control wound condition had high levels of F1 at 1 weeks and 3 weeks with a significant decrease at 3 weeks. There was a low, decreasing number of F2 macrophages in the control wound environment, further suggesting that F2 is associated with fibrotic pathology. Finally, we performed immunofluorescence (IF) for F4/80^+^CD9^+^ on tissue treated with PCL to visualize F2 (and FP1) macrophages (Fig. 5G).

### The pathological F2 macrophage is dependent on IL-17 and is present in human disease

Based on the evidence suggesting F2 is unique to the PCL condition and pathological fibrosis, coupled with the suggested feedback loop between IL-36γ and IL-17, we further evaluated the F2 population and its dependence on IL-17 signaling. PCL was implanted in mice lacking IL-17A (*Il17a*^*-/-*^) or the IL-17 receptor (*Il17ra*^*-/-*^) to ablate IL-17 signaling. Fibrosis in response to PCL decreased in *Il17a*^*-/-*^ mice but was only completely abrogated in *Il17ra*^*-/-*^ mice (*19*). Connecting this functional outcome with macrophage responses, immunostaining for CD9 and F/480 decreased significantly in *Il17a*^*-/-*^ and *Il17ra*^*-/-*^ mice treated with PCL compared to WT after 12 weeks (Fig. 6A). *Il36γ*, coding for a key cytokine expressed in F2 that links IL-17 and autoimmune responses in the tissue, significantly increased after PCL implantation compared to saline in WT animals. Expression of *Il36γ* decreased significantly in *Il17a*^*-/-*^ and *Il17ra*^*-/-*^ mice, further supporting a connection between IL-17 signaling and IL-36*γ* in F2 (Fig. 6B).

**Figure 6.**
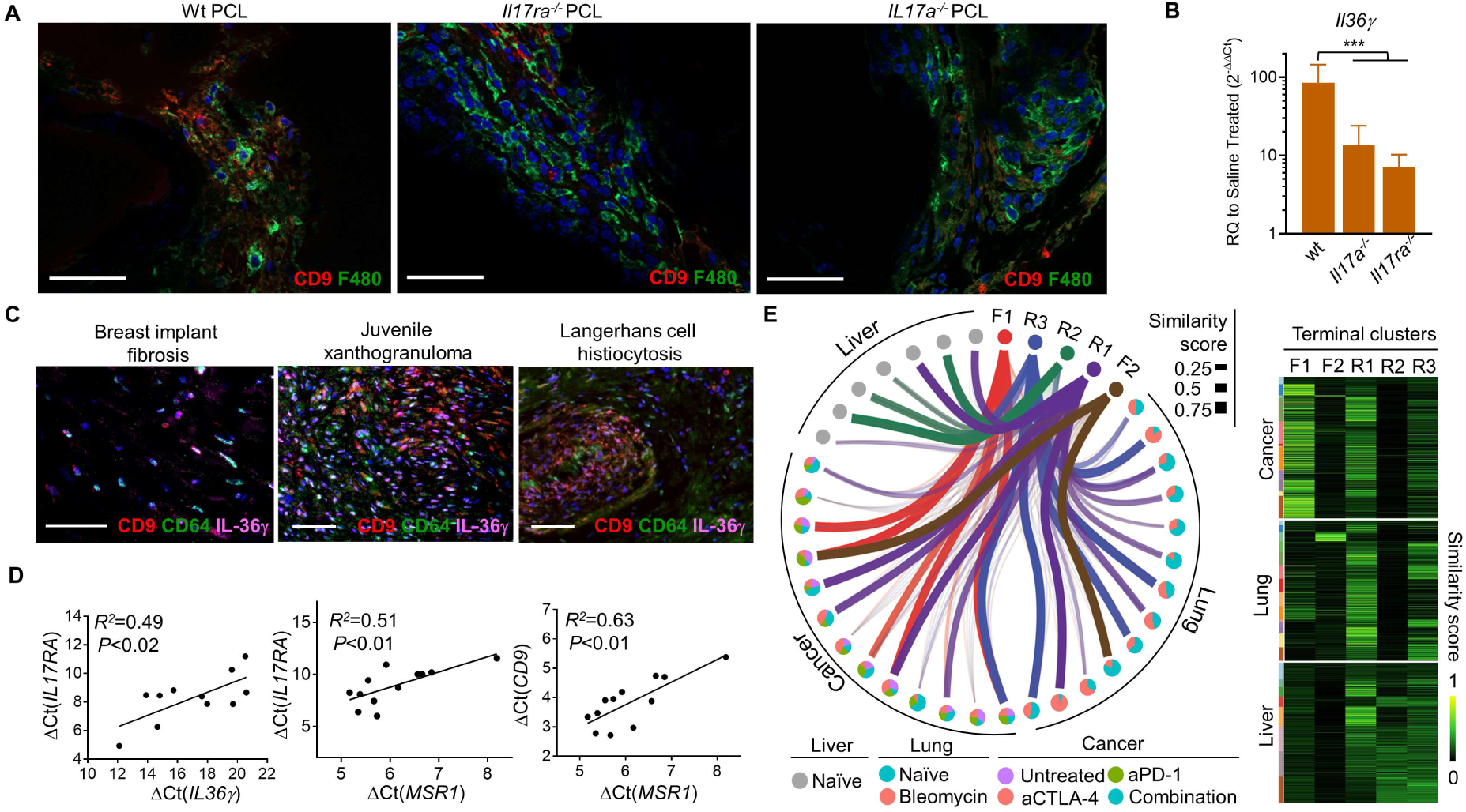
Profibrotic CD9^hi+^*IL-36γ*^+^ macrophages are dependent on IL-17 signaling and terminal clusters are relevant in various pathologies. (A) Immunofluorescent staining for mouse macrophage markers F4/80 and CD9 in wild type, *Il17ra^-/-^*, and *Il17a^-/-^* mice 12 weeks after implantation with PCL (scale bars = 50 μm). (B) *Il36γ* gene expression in wild type, *Il17ra^-/-^*, and *IL17a^-/-^* mice with PCL normalized to saline controls (n=4, biologically independent, ANOVA with multiple comparison, \*\*\**p* <0.001). (C) Immunofluorescent staining for CD64, CD9, IL-36γ positive macrophages in human breast implant tissue capsules (scale bars = 50 μm), juvenile xanthogranuloma, and Langerhans cell histiocytosis (scale bars = 200 μm). (D) Gene expression correlations of human *IL17RA with IL36γ, IL17RA* and *CD9* with *MSR1* in human breast implant fibrotic capsules. (E) Network diagrams and similarity heatmaps for terminal fibrotic and regenerative macrophage clusters to clusters from repository single cell RNA data sets for murine models of cancer (sarcoma +/-immunotherapies aCTLA-4, aPD-1), lung fibrosis (+/- bleomycin induction) and human liver. Circles represent percent compositions of clusters by condition.

The broader relevance of the fibrosis-associated F2 profile was explored in human fibrotic pathologies. Positive immunofluorescence staining for F2-specific markers (CD64^+^CD9^+^IL-36*γ* ^+^) in human breast implant tissue capsules as well as histiocytosis (juvenile xanthogranuloma and Langerhans cell histiocytosis) suggest that these macrophages are also relevant to human disease (Fig. 6C). Further, *IL17RA* expression is correlated to *IL36γ* and the expression of the macrophage marker *MSR1*, supporting a role of the IL-17/IL-36γ feedback loop in human macrophages (Fig.6D).

### Terminal macrophages profiles are present in human and murine tissues

To determine the broader relevance of the model biomaterial-induced macrophages, we utilized the SingleCellNet program (*32*) and trained the scRNAseq cell classification algorithm using the terminal macrophage data sets (Supplementary Fig. 13). We then compared the results to macrophages computationally extracted from publicly available data sets of healthy human liver (*33*), a mouse model of idiopathic pulmonary fibrosis (*34*), and a mouse model of sarcoma with cancer immunotherapy (*15*). After clustering macrophages within each data set, we applied SingleCellNet to quantify similarity with the terminal biomaterial macrophages (Figure 6E). The R1 and R3 macrophage subsets were found in all of the conditions that we evaluated. While the expression of muscle markers in R3 initially suggest that they may be unique to the muscle tissue environment, their presence in all the conditions studied suggest that even this subset has broader relevance in multiple tissue types and pathologies. We found that only liver, a strongly regenerative tissue, contained macrophages similar to R2. The majority of macrophages in the sarcoma were similar to F1 but there was also a small cluster similar to F2. The F2 cluster was also present in the lung tissue.

## Discussion

Macrophages play a critical role in execution of host immune responses to infection, cancer, wound healing, and maintaining tissue homeostasis. Here, we applied single cell RNA sequencing to identify and characterize macrophage phenotypes associated with tissue microenvironments modeled using biomaterials that promote divergent tissue environments, immune profiles, and functional outcomes. The unbiased classification and characterization from the single cell analysis provide new, distinct phenotypic profiles that can be identified using a combination of surface markers and standard experimental flow cytometry and immunostaining techniques. Terminal clusters discovered from single cell analysis provide refined phenotypic and functional macrophage characterization in different tissue environments. We identified a previously unknown macrophage population that links a fibrotic tissue environment with Type 17 immune responses and signatures of autoimmunity, which was abrogated with loss of IL-17 signaling.

The macrophages associated with UBM and IL-4 in the tissue are heterogeneous and distributed in phenotypically distinct clusters. IL-4 is a cytokine recognized for promoting repair of muscle (*4*), liver (*35*)., and cartilage (*36*), and is critical for macrophage polarization in a healing wound (*37*). The UBM environment induced greater macrophage heterogeneity with two primary terminal subsets with phenotypes relevant to tissue repair. Expression analysis of the R1 cluster suggests it is important for mobilizing and educating immune cells through the expression of chemokines and increased antigen presentation that is required during the early wound healing process. The R2 macrophages, with the highest level of *Il4ra*, expressed genes relevant to stimulation of other cell types important for Type 2 responses and regeneration including *Ccl24* (Eotaxin-2), coding for a protein that attracts and activates eosinophils. This finding is supported by Chawla et al., that demonstrated the IL-4 secreting eosinophils are critical to muscle repair (*38*). The metabolic profiles of the R1 (glycolysis) and R2 (oxidative phosphorylation) correlate with distinct functions of antigen presentation and adaptive-related chemokine expression versus phagocytosis, that was validated experimentally in sorted R2 macrophages. Glycolysis has been associated with inflammatory macrophages (*39*) and oxidative phosphorylation with alternatively activated macrophages but macrophages (*40*). *In vitro* studies of conventional M2 macrophages required inhibition of both metabolic pathways to inhibit IL-4 induced STAT6 phosphorylation (*41*).

The distribution of macrophages isolated from the PCL-treated wounds was less heterogeneous than the ECM-treated tissues and included the functional subsets F1 and F2. The F1 cluster expresses many genes associated with inflammation including interferon-related cytokines and activation of the innate and adaptive immune system. The R1 cluster also expressed markers of inflammation and mobilization but the magnitude of expression and types of inflammatory markers were significantly different. This difference in the F1 and R1 inflammatory profile suggests the importance of the early inflammatory response in directing the subsequent tissue repair or development of a foreign body capsule or fibrosis. The time course of flow cytometry revealed that R1 increased with ECM treatment. Since ECM treatment improves tissue repair, R1 may represent an inflammatory phenotype that can be targeted to enhance tissue development.

The F2 cluster associated with PCL treatment expressed genes that connected Type 17 immunity and markers of autoimmune disease. Type 17 immune responses are associated with autoimmunity in diseases such as psoriasis, irritable bowel syndrome and inflammatory arthritis (*28, 42-44*). The F2 macrophages express IL-36γ, a cytokine that is found clinically in the skin of psoriasis patients and in inflammatory arthritis (*45*). This cytokine was also recently identified as a target in tumors that, when blocked, enhances responsiveness to immunotherapy (*46*). It is also implicated in a positive feedback loop with IL-17 (*28*). In other work, we demonstrated that IL-17 is produced by innate lymphocytes, γδ and CD4^+^ T cells in response to PCL implantation in mice and in the fibrous capsule surrounding human breast implants (*19*). IL-17 is implicated in fibrotic disease in in the lung (*30*), heart (*31*), and liver (*47*) in addition to the foreign body response (*5*). The F2 macrophage population expressing IL-17 receptor A, links n IL-17 signaling, fibrosis and autoimmune disease.

The F2 macrophages also expressed multiple forms of *Trem* (triggering receptor expressed on myeloid cells) and its ligands that are associated with autoimmune diseases such as inflammatory bowel disease and psoriasis (*48, 49*). TREM integrates and broadly modifies inflammatory signals across the innate and adaptive immune system. The presence of F2 surface markers and related cytokines in human tissues provides evidence that the tissue immune environment created by PCL and the mechanisms of response may be broadly relevant to various pathological conditions. Further supporting the broader relevance of the macrophage subsets, we found macrophages clusters similar to F2 in publicly available data sets of idiopathic lung fibrosis and sarcoma. Multiple genes in the F2 subset have functions that remain unknown. Based on the potential importance of this subset in disease pathology, further studies into these unknown genes may be warranted. While small in number, the F2 macrophage subset, is highly differentiated and may play a critical role in pathologies associated with the pro-inflammatory macrophages in tissue fibrosis. Further studies to investigate the macrophage subsets that we identified by single cell analysis will determine if the surface markers and their associated subpopulations and respective expression profiles are stable and broadly relevant. The functional impact of removing or augmenting specific macrophage subsets will provide further insight into their mechanistic contributions to the tissue environment across multiple pathologies.

## Acknowledgements

We gratefully acknowledge funding from the Morton Goldberg Chair, the Bloomberg∼Kimmel Institute for Cancer Immunotherapy, and Acell Inc. We would like to thank the Cindy Sears laboratory for providing the original breeding mice strain for transgenic *Il17ra*^*-/-*^, *Il17a*^*-/-*^.

## Author contributions

S.D.S., C.C., J.H.E. conceptualized and drafted figures and manuscript. S.D.S., C.C., J.H.E., D.M.P., P.C., F.H. contributed to experimental design and interpretation of results. S.D.S., C.C., L.C., D.R.M, R.S., A.T., J.S., J.T. contributed to conducting experimental procedures and analyzing data. All authors participated in editing and revising the manuscript text and figures.

## Methods

Contact for Reagent and Resource Sharing

Requests regarding protocols and resources should be directed to and will be fulfilled by the lead contact J Elisseeff

## References

1. S. Aras, M. R. Zaidi, TAMeless traitors: macrophages in cancer progression and metastasis. British journal of cancer 117, 1583 (2017).

2. V. Kumar, S. Patel, E. Tcyganov, D. I. Gabrilovich, The nature of myeloid-derived suppressor cells in the tumor microenvironment. Trends in immunology 37, 208–220 (2016).

3. S. F. Badylak, J. E. Valentin, A. K. Ravindra, G. P. McCabe, A. M. Stewart-Akers, Macrophage phenotype as a determinant of biologic scaffold remodeling. Tissue Engineering Part A 14, 1835–1842 (2008).

4. K. Sadtler et al., Developing a pro-regenerative biomaterial scaffold microenvironment requires T helper 2 cells. Science 352, 366–370 (2016).

5. T. A. Wynn, K. M. Vannella, Macrophages in tissue repair, regeneration, and fibrosis. Immunity 44, 450–462 (2016).

6. Q. N. Myrvik, E. S. Leake, B. Fariss, Studies on pulmonary alveolar macrophages from the normal rabbit: a technique to procure them in a high state of purity. The Journal of Immunology 86, 128–132 (1961).

7. D. L. Knook, N. Blansjaar, E. C. Sleyster, Isolation and characterization of Kupffer and endothelial cells from the rat liver. Experimental cell research 109, 317–329 (1977).

8. G. J. Guillemin, B. J. Brew, Microglia, macrophages, perivascular macrophages, and pericytes: a review of function and identification. Journal of leukocyte biology 75, 388–397 (2004).

9. Y. Lavin et al., Tissue-resident macrophage enhancer landscapes are shaped by the local microenvironment. Cell 159, 1312–1326 (2014).

10. P. J. Murray, T. A. Wynn, Protective and pathogenic functions of macrophage subsets. Nature reviews immunology 11, 723 (2011).

11. A. Mantovani et al., The chemokine system in diverse forms of macrophage activation and polarization. Trends in immunology 25, 677–686 (2004).

12. C. D. Mills, K. Kincaid, J. M. Alt, M. J. Heilman, A. M. Hill, M-1/M-2 macrophages and the Th1/Th2 paradigm. The Journal of Immunology 164, 6166–6173 (2000).

13. D. M. Mosser, J. P. Edwards, Exploring the full spectrum of macrophage activation. Nature reviews immunology 8, 958 (2008).

14. P. J. Murray et al., Macrophage activation and polarization: nomenclature and experimental guidelines. Immunity 41, 14–20 (2014).

15. M. M. Gubin et al., High-dimensional analysis delineates myeloid and lymphoid compartment remodeling during successful immune-checkpoint cancer therapy. Cell 175, 1014–1030 (2018).

16. A.-C. Villani et al., Single-cell RNA-seq reveals new types of human blood dendritic cells, monocytes, and progenitors. Science 356, eaah4573 (2017).

17. J. M. Anderson, A. Rodriguez, D. T. Chang. Foreign body reaction to biomaterials, in Seminars in immunology, (Elsevier), vol. 20, pp. 86–100.

18. B. D. Ratner, A. S. Hoffman, F. J. Schoen, J. E. Lemons, Biomaterials science: an introduction to materials in medicine. (Elsevier, 2004).

19. L. Chung et al., Interleukin-17 and senescence regulate the foreign body response. bioRxiv, 583757 (2019).

20. B. M. Sicari et al., An acellular biologic scaffold promotes skeletal muscle formation in mice and humans with volumetric muscle loss. Science translational medicine 6, 234ra258–234ra258 (2014).

21. H. Kimmel, M. Rahn, T. W. Gilbert, The clinical effectiveness in wound healing with extracellular matrix derived from porcine urinary bladder matrix: a case series on severe chronic wounds. The Journal of the American College of Certified Wound Specialists 2, 55–59 (2010).

22. J. Zografakis et al., Urinary bladder matrix reinforcement for laparoscopic hiatal hernia repair. JSLS: Journal of the Society of Laparoendoscopic Surgeons 22, (2018).

23. J. C. Doloff et al., Colony stimulating factor-1 receptor is a central component of the foreign body response to biomaterial implants in rodents and non-human primates. Nature materials 16, 671 (2017).

24. E. L. Gautier et al., Gene-expression profiles and transcriptional regulatory pathways that underlie the identity and diversity of mouse tissue macrophages. Nature immunology 13, 1118 (2012).

25. M. T. Wolf et al., A biologic scaffold–associated type 2 immune microenvironment inhibits tumor formation and synergizes with checkpoint immunotherapy. Science translational medicine 11, eaat7973 (2019).

26. K. Sadtler et al., Divergent immune responses to synthetic and biological scaffolds. Biomaterials 192, 405–415 (2019).

27. C. J. Workman et al., LAG-3 regulates plasmacytoid dendritic cell homeostasis. The Journal of Immunology 182, 1885–1891 (2009).

28. Y. Carrier et al., Inter-regulation of Th17 cytokines and the IL-36 cytokines in vitro and in vivo: implications in psoriasis pathogenesis. Journal of Investigative Dermatology 131, 2428–2437 (2011).

29. L. J. Celada et al., PD-1 up-regulation on CD4+ T cells promotes pulmonary fibrosis through STAT3-mediated IL-17A and TGF-β1 production. Science translational medicine 10, eaar8356 (2018).

30. M. S. Wilson et al., Bleomycin and IL-1β–mediated pulmonary fibrosis is IL-17A dependent. Journal of Experimental Medicine 207, 535–552 (2010).

31. L. Wu et al., Cardiac fibroblasts mediate IL-17A–driven inflammatory dilated cardiomyopathy. Journal of Experimental Medicine 211, 1449–1464 (2014).

32. Y. Tan, P. Cahan, SingleCellNet: a computational tool to classify single cell RNA-Seq data across platforms and across species. bioRxiv, 508085 (2018).

33. S. A. MacParland et al., Single cell RNA sequencing of human liver reveals distinct intrahepatic macrophage populations. Nature communications 9, 4383 (2018).

34. D. Aran et al., Reference-based analysis of lung single-cell sequencing reveals a transitional profibrotic macrophage. Nature immunology, 1 (2019).

35. C. Blériot et al., Liver-resident macrophage necroptosis orchestrates type 1 microbicidal inflammation and type-2-mediated tissue repair during bacterial infection. Immunity 42, 145–158 (2015).

36. M. F. Pustjens et al., IL4-10 synerkine induces direct and indirect structural cartilage repair in osteoarthritis. Osteoarthritis and Cartilage 24, S532 (2016).

37. K. Sadtler et al., The scaffold immune microenvironment: biomaterial-mediated immune polarization in traumatic and nontraumatic applications. Tissue Engineering Part A 23, 1044–1053 (2017).

38. J. E. Heredia et al., Type 2 innate signals stimulate fibro/adipogenic progenitors to facilitate muscle regeneration. Cell 153, 376–388 (2013).

39. J.-C. Rodríguez-Prados et al., Substrate fate in activated macrophages: a comparison between innate, classic, and alternative activation. The Journal of Immunology 185, 605–614 (2010).

40. J. I. Odegaard, A. Chawla, Alternative macrophage activation and metabolism. Annual Review of Pathology: Mechanisms of Disease 6, 275–297 (2011).

41. F. Wang et al., Glycolytic stimulation is not a requirement for M2 macrophage differentiation. Cell metabolism 28, 463–475 (2018).

42. G. Fiorino, P. D. Omodei, Psoriasis and Inflammatory Bowel Disease: Two Sides of the Same Coin? Journal of Crohn’s and Colitis 9, 697–698 (2015).

43. S. Kagami, H. L. Rizzo, J. J. Lee, Y. Koguchi, A. Blauvelt, Circulating Th17, Th22, and Th1 cells are increased in psoriasis. Journal of Investigative Dermatology 130, 1373–1383 (2010).

44. J. F. Zambrano-Zaragoza, E. J. Romo-Martínez, M. Durán-Avelar, N. García-Magallanes, N. Vibanco-Pérez, Th17 cells in autoimmune and infectious diseases. International journal of inflammation 2014, (2014).

45. M. T. Mizwicki et al., Tocilizumab attenuates inflammation in ALS patients through inhibition of IL6 receptor signaling. American journal of neurodegenerative disease 1, 305 (2012).

46. X. Wang et al., IL-36γ transforms the tumor microenvironment and promotes type 1 lymphocyte-mediated antitumor immune responses. Cancer cell 28, 296–306 (2015).

47. W. Seo et al., Exosome-mediated activation of toll-like receptor 3 in stellate cells stimulates interleukin-17 production by γδ T cells in liver fibrosis. Hepatology 64, 616–631 (2016).

48. J. W. Ford, D. W. McVicar, TREM and TREM-like receptors in inflammation and disease. Current opinion in immunology 21, 38–46 (2009).

49. M. Schenk, A. Bouchon, F. Seibold, C. Mueller, TREM-1–expressing intestinal macrophages crucially amplify chronic inflammation in experimental colitis and inflammatory bowel diseases. The Journal of clinical investigation 117, 3097–3106 (2007).

50. K. J. Livak, T. D. Schmittgen, Analysis of relative gene expression data using real-time quantitative PCR and the 2-ΔΔCT method. methods 25, 402–408 (2001).

51. A. Dobin et al., STAR: ultrafast universal RNA-seq aligner. Bioinformatics 29, 15–21 (2013).

52. A. Butler, P. Hoffman, P. Smibert, E. Papalexi, R. Satija, Integrating single-cell transcriptomic data across different conditions, technologies, and species. Nature biotechnology 36, 411 (2018).

53. M. S. Kowalczyk et al., Single-cell RNA-seq reveals changes in cell cycle and differentiation programs upon aging of hematopoietic stem cells. Genome research 25, 1860–1872 (2015).

54. V. Y. Kiselev et al., SC3: consensus clustering of single-cell RNA-seq data. Nature methods 14, 483 (2017).

55. M. D. Robinson, D. J. McCarthy, G. K. Smyth, edgeR: a Bioconductor package for differential expression analysis of digital gene expression data. Bioinformatics 26, 139–140 (2010).

56. A. Sergushichev, An algorithm for fast preranked gene set enrichment analysis using cumulative statistic calculation. BioRxiv, 060012 (2016).

57. D. Szklarczyk et al., STRING v10: protein–protein interaction networks, integrated over the tree of life. Nucleic acids research 43, D447–D452 (2014).

58. C. Von Mering et al., STRING: known and predicted protein–protein associations, integrated and transferred across organisms. Nucleic acids research 33, D433–D437 (2005).

59. M. Bastian, S. Heymann, M. Jacomy, Gephi: an open source software for exploring and manipulating networks, in Third International AAAI Conference. (2009).

60. K. Street et al., Slingshot: Cell lineage and pseudotime inference for single-cell transcriptomics. BMC genomics 19, 477 (2018).

61. G. La Manno et al., RNA velocity of single cells. Nature 560, 494 (2018).

62. C. Tabula Muris, Single-cell transcriptomics of 20 mouse organs creates a Tabula Muris. Nature 562, 367 (2018).

