## Supplementary materials for "Single cell RNA-seq in regenerative and fibrotic biomaterial environments defines new macrophage subsets"

##### Methods

Contact for Reagent and Resource Sharing

##### EXPERIMENTAL MODEL AND SUBJECT DETAILS

*Surgical procedures and implantation*
All animal procedures were performed in adherence to approved JHU IACUC protocols. Animals included female mice wild type C57BL/6j (Jackson Laboratories) and transgenic *Il17a*$^{-/-}$ (Dr. Yoichiro Iwakura, University of Tokyo, Tokyo, Japan) and *IL17ra*$^{-/-}$ (Dr. Tomas Mustelin, Amgen, Seattle) at ages from 8-10 weeks. The bilateral traumatic muscle defect was created as previously described (*4*). The defects were filled with 30 mg of a synthetic material or biological scaffold material. PCL was employed as a synthetic material (particulate, $M_n$ = 50,000 g/mol, mean particle size < 600 µm, Polysciences). In turn, as a biological scaffold material, decellularized urinary bladder matrix (Matristem, Acell) was implanted from 0.05 ml of a 400 mg/ml suspension in phosphate buffered saline (PBS). Control surgeries were injected with 0.05 ml of PBS as a no implant control. All materials were UV sterilized prior to use. Mice were given subcutaneous carprofen (Rimadyl®, Zoetis) at 5 mg/kg for pain relief. For the sample harvest, mice were euthanized at 1, 3, 6, 12-weeks post-surgery.

*Specimen harvest*
We obtained murine samples by dissecting the quadriceps femoris muscle followed by fine dicing. Tissues were digested for 45 min at 37°C with 1.67 Wünsch U/ml Liberase TL (Roche Diagnostics) and 0.2 mg/ml DNase I (Roche Diagnostics) in RPMI 1640 medium (Gibco). The digested tissues were ground through 70 µm cell strainers (ThermoFisher Scientific) and rinsed with PBS + 0.05% bovine serum albumin and then washed twice with 1X PBS. The enriched single cell suspension was washed and stained with the following antibody panels, respective to the application.

*Flow cytometry and fluorescence activated cell-sorting*
For cell isolation using FACS, suspensions of single cells from digested muscles were stained for 20 minutes at 4°C using Viability Dye eFluor™ 780 (eBioscience). For single cell analysis, staining in an antibody panel was conducted 30 minutes at 4°C including F4/80 PE-Cy7 (BioLegend), CD11b AlexaFluor700 (BioLegend), CD64 PerCP-Cy5.5 (BioLegend), MHCII (I-A/I-E) AlexaFluor488 (BioLegend), CD3 APC (BioLegend), Ly6c BrilliantViolet510 (BioLegend), CD45 BrilliantViolet605 (BioLegend), and Fc Block TruStain fcX (anti-mouse CD16/32) (BioLegend). Sorting of murine macrophages (CD45$^{+}$F4/80$^{hi+}$CD64$^{+}$) was performed from live,CD45$^{+}$CD3$^{-}$ using a BD FACSAria II (BD Biosciences). The post-sort purity of macrophages used for single cell RNA sequencing was > 98%. For subset analysis, an Attune NxT Flow Cytometer (ThermoFisher Scientific) after viability stain, a panel comprised of F4/80 PE-Cy7 (BioLegend), CD9 FITC (BioLegend), CD11c AlexaFluor700 (BioLegend), MHCII (I-A/I-E) PE-CF594 (eBiosciences), CD301b PE (BioLegend), CD206 APC (BioLegend), CD86 BrilliantViolet410 (BioLegend), Ly6c BrilliantViolet510 (BioLegend), CD45 BrilliantViolet605 (BioLegend), Fc Block TruStain FcX (anti-mouse CD16/32) (BioLegend). Data analysis was performed in FlowJo Flow Cytometry Analysis Software (Treestar). For tSNE projection of multi-dimensional flow cytometry datasets, downsampling of Macrophages (CD45$^{+}$F4/80$^{hi+}$) were performed to 50,400 cells total of samples from regenerative, fibrotic, saline control conditions

(n=4, biologically independent). Then samples were concatenated and tSNE projection was computed (Iterations = 1000, Perplexity = 30) (Supplementary Fig. 14). For visualization of separation macrophage subtypes were back gated onto 2-dimensional projection.

Macrophage FACS panel:

| Fluorophore | Marker | Manufacturer |
|---|---|---|
| eFluor780 | Live/Dead | Life Technologies |
| AF700 | CD11b | eBioscience |
| PerCP-Cy5.5 | CD64 | BioLegend |
| PE-Cy7 | F4/80 | BioLegend |
| APC | CD3 | BioLegend |
| AF488 | MHCII | BioLegend |
| BV510 | Ly6c | BioLegend |
| BV605 | CD45 | BioLegend |

Macrophage Subtype panel:

| Fluorophore | Marker | Manufacturer |
|---|---|---|
| eFluor780 | Live/Dead | Life Technologies |
| FITC | CD9 | BioLegend |
| PE | CD301b | BioLegend |
| PE-CF594 | MHCII | BioLegend |
| PE-Cy7 | F4/80 | BioLegend |
| APC | CD206 | BioLegend |
| AF700 | CD11c | BioLegend |
| BV421 | CD86 | BioLegend |
| BV510 | Ly6c | BioLegend |
| BV605 | CD45 | BioLegend |

*Histopathology and Immunofluorescence microscopy*

After harvest, implanted tissues at 1, 3, 6, and 12 weeks after implantation were fixed in 10% neutral buffered formalin for 48 hours before ethanol and xylenes dehydration. Archival formalin-fixed paraffin-embedded tissues from patients with conditions known to involve increased tissue macrophages, such as Langerhans cell histiocytosis, juvenile xanthogranuloma, and dermal scar, were obtained from the Johns Hopkins Hospital surgical pathology archives. Human tissue samples including silicone breast implants were acquired from patients undergoing implant exchange or replacement surgeries exemption IRB00088842. The average age was 56 years old (range of 41-70 years old) and average implant residence time was 41 months (range of 1-360 months). Paraffin-embedded sections of 7 μm thickness were produced using a microtome (Leica RM2255 microtome). Immunofluorescence staining for CD9, IL-36γ, F4/80, CD301b was performed with a tyramide signal amplification (TSA) method using consecutive staining with Opal-520, Opal-570, and Opal-650 respectively. Histological slides were rehydrated in an ethanol to water gradient. After post-fixation in 10% neutral buffered formalin for 30 mins, antigen retrieval was conducted in 1 x AR6 buffer (Perkin-Elmer) at 95°C for 15 min. After cooling, endogenous peroxidases were quenched in 3% (v/v) aqueous peroxide (Sigma-Aldrich.) All staining was conducted at room temperature after blocking with antibody block/diluent (Perkin-Elmer). For each staining round, the primary antibody in block/diluent (Perkin-Elmer) was incubated at room temperature for 60 mins, followed by 10 mins of incubation with HRP polymer conjugated secondary antibody, and 10 mins of Opal reagent (1:150) in 1 x plus amplification diluent (Perkin-Elmer). The Antibody HRP complex was stripped in a microwave at 95°C 1 x AR6 buffer (Perkin-Elmer) for 15 mins. After quenching of autofluorescence in 0.04% (w/v) of Sudan Black (Sigma-Aldrich) dissolved in 70%(v/v) ethanol, slides were then counterstained with DAPI for 5 mins before being mounted using DAKO mounting medium. Imaging of the histological samples was performed on a Zeiss Axio Imager A2 and Zeiss AxioVision software ver. 4.2.

*Phagocytosis assay*

Macrophage subsets R1, R2, were sorted according to the gating strategy described in Supplementary Figure 4. Around 100,000 cells were collected in RPMI + 5% (v/v) fetal bovine serum (Gibco). Macrophages were seeded at 60,000 cells/cm$^2$ and incubated at 37°C for 12 hours. Then phagocytosis was conducted for 4 hours at a concentration of 1.0 x 10^6 cells/ml with Fluoresbrite flash red (Polysciences) with an average diameter of 2.00 μm particles at 2.0 x 10^8 particles/ml. As readout to quantify macrophage subset phagocytosis fluorescence microscopy using Alexa Fluor 594 phalloidin (Invitrogen) and DAPI.

*qRT-PCR gene expression assay*

Total and enriched mRNA was isolated from whole tissue using TRIzol reagent and Qiagen's RNeasy kits. All qRT-PCR was performed using TaqMan Gene Expression Master Mix (Applied Biosystems) according to manufacturer's instructions. Briefly, 2 μg of enriched mRNA was used to synthesize cDNA using Superscript IV VILO Master Mix (ThermoFisher Scientific). The cDNA concentration was set to 50 - 100 ng/well (in a total volume of 20 μL PCR reaction) to match manufacturer recommendations. The qRT-PCR reactions were performed on the StepOne Plus Real-Time PCR System (ThermoFisher Scientific), as TaqMan single-plex assays, using manufacturer recommended settings for quantitative and relative expression. For tissue samples, β2m was used as the reference gene and samples were normalized to PBS treated controls. All qRT-PCR data was analyzed using the Livak Method, wherein ΔΔCt values are calculated and reported as relative quantification values (RQ) calculated by $2^{-\Delta\Delta Ct}$ (*50*). RQ values are represented by the geometric means with error bars representing the geometric standard deviation. All qRT-PCR assays were completed within the laboratory by the authors involved with the study. Low expressing mRNA transcripts were pre-amplified using the TaqMan Pre-Amp system according to manufacturer recommendations with 14 cycles of amplification with the primer probes of interest.

Murine TaqMan Gene Expression assay Probes:

| Primer | Assay ID: |
|---|---|
| *β2m* | Mm00437762 |
| *Il36γ* | Mm00463327 |

Human TaqMan Gene Expression assay Probes:

| Primer | Assay ID: |
|---|---|
| *β2M* | Hs00187842_m1 |
| *IL36γ* | Hs00219742_m1 |
| *MSR1* | Hs00234007_m1 |
| *CD9* | Hs01124026_m1 |
| *IL17RA* | Hs01056316_m1 |

*Single cell encapsulation and Library generation*
After sorting of Macrophages ($CD45^{+}F4/80^{hi+}$), single cells were encapsulated in water-in-oil emulsion along with gel beads coated with unique molecular barcodes using the 10x Genomics Chromium Single-Cell Platform. For single cell RNA library generation, the manufacturers' protocol was performed (10X Single Cell 3' v2). Sequencing was performed using an Illumina HiSeq2500 Rapid Mode with 310 million reads per sample and a sequencing configuration of 26x8x98 (UMI x Index x Transcript read). The Cell Ranger pipeline software was used to align reads and generate expression matrices for downstream analysis.

**COMPUTATIONAL ANALYSIS**
*Sequence alignment, filtering, normalization, and scaling*
Alignment was performed with STAR (*51*) through the Cell Ranger pipeline. Filtering, normalization, and scaling were performed using Seurat (*16, 52*). Cells with UMI counts for fewer than 200 genes and genes with expression in less than 0.1% of cells were both dropped from analysis. Data was normalized by $E_{norm}=\log(UMI*10{,}000/UMI_{total})$ where $UMI_{total}$ is total UMI expression for a given gene. Scaling was performed to remove unwanted effects correlated to batch, total UMI count, percent of mitochondrial genes, and cell cycle. To determine cell cycle scores for use with scaling, Seurat's CellCycleScoring function was used with a previously determined set of genes (*53*) correlated with G2M or S phase. Data scaling was performed by first fitting a linear model with the parameters to scale out as independent variables (batch, total UMI count, percent mitochondrial genes, G2M score, and S score) (Supplementary Fig. 15). For each gene, the residuals from the fit were Z-scored and used for downstream analysis. Finally, principal components were calculated and the top 50 were selected based on leveling of variance per principal component as determined by an elbow plot (Supplementary Fig. 16).

*Clustering analysis*
SC3 (*54*) and Seurat's unsupervised clustering algorithms using the top 50 principal components were compared using silhouette coefficients. For each clustering algorithm, a reasonable range of resolution or k parameters were determined *a priori*. Silhouette coefficients were determined for each result and compared. Seurat was found to have consistently higher silhouette coefficients and was used. A resolution value was selected based on a combination of

high silhouette coefficient and reasonable number of clusters for biological consideration. After clustering, a small population of contaminant fibroblasts were found and removed (Supplementary Fig. 17).

*Removal of low signal clusters*
Downstream analysis also found evidence of clusters which had little signal remaining after scaling. To attempt to quantify this, differential expression test was calculated using Seurat's negative binomial statistical test with a minimum log fold-change of 1. Based on these results together with quality control information such as total UMI count (Supplementary Fig. 2), we decided to remove clusters O1-4 from analysis to avoid diluting the signal from clusters with stronger signal. The resulting data matrix was reprocessed identically to ultimately find 9 clusters of macrophages with strong enough signal to identify surface markers and suggest biological function.

*Differential expression and gene set enrichment*
Differential expression was calculated using edgeR's exactTest (*55*). Unless otherwise specified, differential expression for a specific cluster was determined by comparison against all other clusters. For heatmaps comparing expression of multiple different genes by cluster, gene expression values were normalized against the highest expression value for each given gene. Feature plots and violin plots for gene expression were generated with Seurat's FeaturePlot and VlnPlot respectively using log-normalized expression values. Ranked gene lists were used to calculate gene set enrichment scores GSEA was performed as previously described using fgsea (*56*). Briefly, all genes in the analysis were ranked by p-value with the exception of the direct comparison between R1 and R2 where genes were ranked by log fold-change. For a given gene set, a running enrichment score is calculated by stepping through the ranked gene list and adding to the score when encountering a gene in the gene set or subtracting from the score otherwise. The enrichment score for a gene set is maximum (or minimum if negative) of the running enrichment score with a higher enrichment score indicating higher concentration of genes in the gene set towards the beginning of the ranked gene list. Finally, a null distribution for the gene set is calculated by repeatedly permuting the ranked genes and used to calculate a p-value. Gene ontology cellular components, biological process, and hallmark gene sets were obtained from the Broad Institute for GSEA.

*Protein network analysis*
Functional network analysis of gene products in macrophage clusters was conducted using a databank-based query of the STRING consortium (*57, 58*). Three letter codes of the top 300 differentially expressed genes per cluster were mapped to the *Mus musculus* protein products. Connectivity of proteins in the network was scored ranked from 0 to 1, with 1 being the highest possible confidence weighed meta-data comparison based on experimental protein-protein interaction evidence, Pubmed text mining and curated databases. Gephi network visualization tool was used to plot the cluster protein networks (*59*). Network nodes represent proteins with the size proportional to the number of interactions. Edges represent protein-protein associations with the thickness proportional to the strength of the meta-evidence supporting the interaction.

*Pseudotime analysis*
Pseudotime analysis was performed using Slingshot (*60*). Precursor clusters were determined using differential expression and gene set enrichment for clusters. In particular, we found that precursor clusters shared gene set enrichment patterns (Supplementary Fig. 3). R3 was removed as an outlier due to highly unique gene expression. To determine genes associated with pseudotime progression, genes were regressed on pseudotime as determined by Slingshot using a general additive model with the gam R package. The top 10 genes by p-value were then

used to generate a heatmap, ordering cells by pseudotime. Z-scored scaled residuals expression values were used for the heatmaps.

*RNA velocity analysis*

RNA velocity analysis was performed as previously described (*61*) using the velocyto.py python package for annotating transcripts as spliced or unspliced followed by the velocyto.R R package to perform velocity estimation. Briefly, transcripts are marked as either spliced or unspliced based on the presence or absence of intronic regions in the transcript. For each gene, a simple model of RNA dynamics is then fit to the data. Finally, the RNA velocity is estimated for each cell by looking for over or underrepresentation of spliced to unspliced ratios. RNA velocity is visualized on a UMAP plot with vector fields representing the averaged velocity of nearby cells.

*Analysis of publicly available data sets*

Publicly available scRNAseq data sets were obtained from GEO (GSE115469 (*33*), GSE111664 (*34*), GSE119352 (*15*)). For each data set, we scaled the data and then used SingleCellNet (*32*) trained on the tabula muris data set (*62*) to quantify similarity to known cell types. We then selected only macrophages and alveolar macrophages and completed analysis up through clustering. After determination of optimal clustering resolution, we applied the SingleCellNet program to quantify similarity to the biomaterial-wound macrophages, training the algorithm on the terminal clusters R1, R2, R3, F1, and F2. This generated five similarity scores for each public data set macrophage, one for each terminal cluster. Finally, we calculated cluster level similarity scores by taking the mean of all cells in the cluster.

### Supplementary Figures

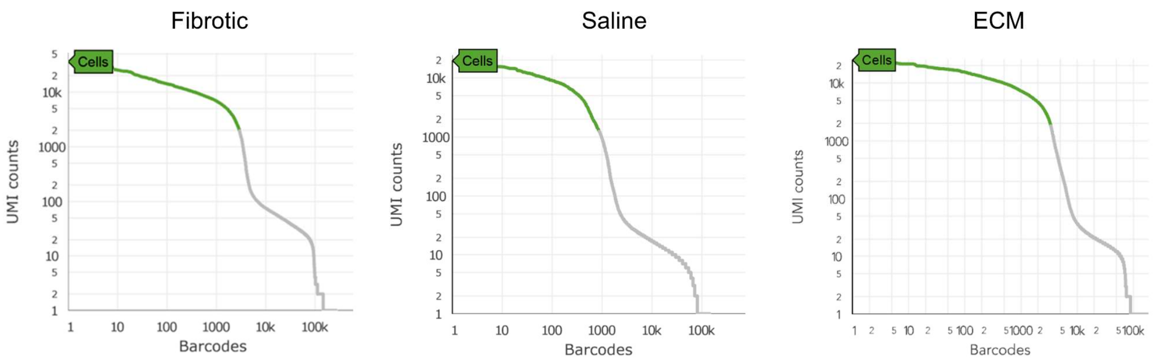


Mean reads per cell: 97594
Median genes per cell: 1906
Total cells after filtering: 2919
Total read no.: 286M
Valid barcodes: 97.1%
>92.5% Q30 bases across all bases

Mean reads per cell: 56825
Median genes per cell: 1528
Total cells after filtering: 876
Total read no.: 286M
Valid barcodes: 97.1%
>92.5% Q30 bases across all bases

Mean reads per cell: 91454
Median genes per cell: 1741
Total cells after filtering: 3343
Total read no.: 286M
Valid barcodes: 97.1%
>92.5% Q30 bases across all bases

**Supplementary Figure 1. Quality control of single cell RNA sequencing data.** Number of UMI counts per barcodes identified in CellRanger software. Green coloring indicates barcodes assigned to cells by CellRanger's algorithms above UMI cutoff minimum and maximum removing debris and doublets.

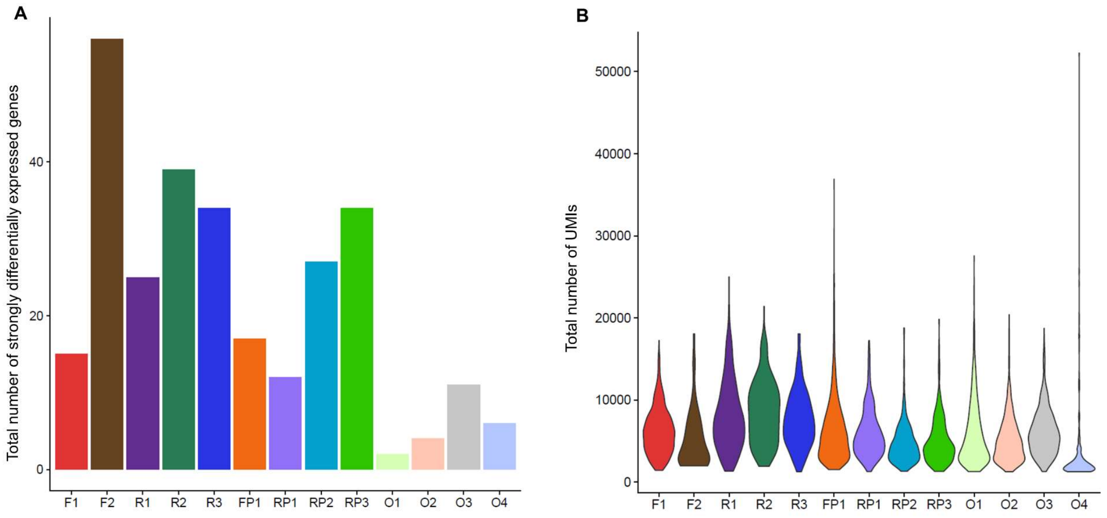


**Supplementary Figure 2. Differential expression and quality control metrics show clusters grouped on non-desirable traits.** (A) Total number of clusters differentially expressed genes with average log fold change greater than 1 or less than -1 are shown. (B) Violin plots by cluster of total UMI counts, a quality control metric related to depth of reads, shows O4 as a cluster of cells with low mean UMI count and high standard deviation.

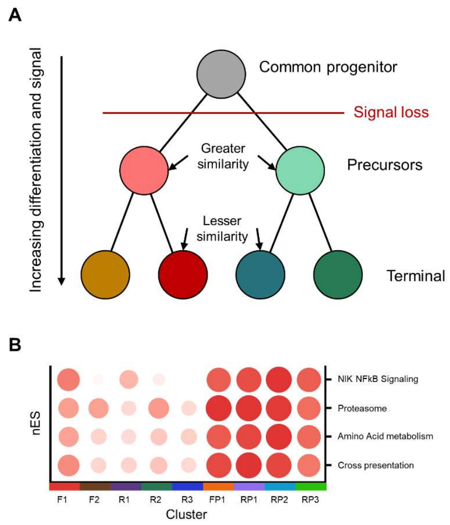


**Supplementary Figure 3. Precursor clusters show similarities across condition.** (A) Model depicting progression of cell differentiation in Slingshot pseudotime analysis by similarity from assumed common progenitor, precursor clusters to terminal clusters. (B) Heatmap of normalized gene set enrichment scores for clusters. Precursor clusters share common gene set enrichment patterns, indicating higher similarity among precursor clusters as compared to terminal clusters.

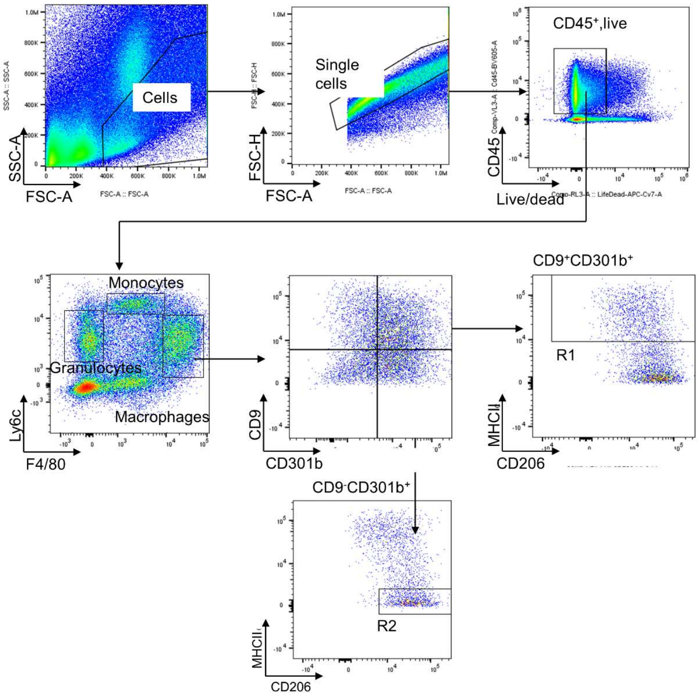


**Supplementary Figure 4. Macrophage gating scheme for regenerative subsets.** Depicted is the multicolor flow cytometry gating strategy to identify subsets R1 ($CD9^{+}CD301^{+}MHCII^{hi+}$) and R2 ($CD9^{+}CD301^{+}$) from single cell suspension of digested murine tissue samples.

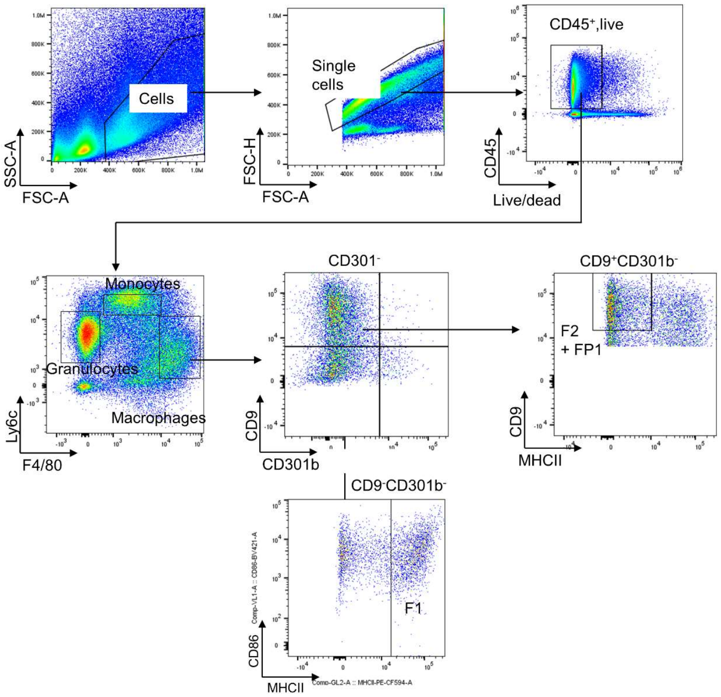


**Supplementary Figure 5. Macrophage gating scheme for fibrotic subsets.** Depicted is the multicolor flow cytometry gating strategy to isolate subsets F1 ($CD9^{-}CD301^{-}MHCII^{hi+}$) and F2 + FP1($CD9^{hi+}CD301^{-}$) from single cell suspension of digested murine tissue samples.

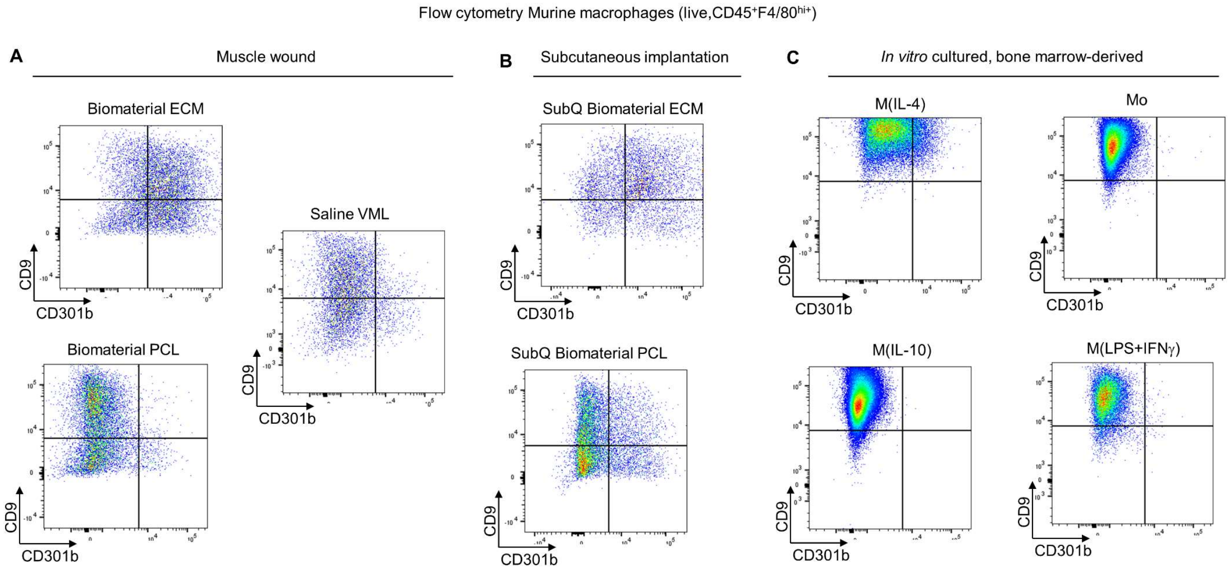


**Supplementary Figure 6. Murine macrophage CD9 and CD301b profiles using flow cytometry.** (A) Volumetric muscle loss model with ECM (regenerative) PCL (fibrotic), saline control. (B) Subcutaneous implantation of ECM and PCL. (C) Bone-marrow derived macrophages cultured on (poly)styrene, differentiated with M-CSF (100 ng/ml)for 1 week, then chemically polarized for 2 days in Ham's F12 + 10% FBS + 1% P/S supplemented with either IL-4 (20 ng/ml), IFNγ (20 ng/ml) + LPS (100 ng/ml), IL-10 (20 ng/ml), or control without stimulation.

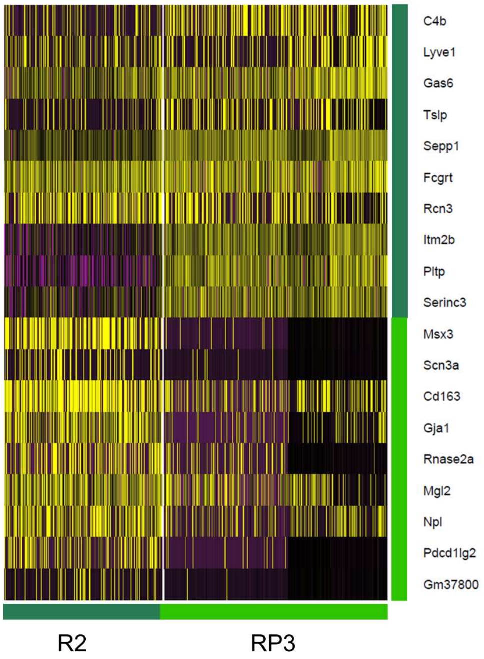


**Supplementary Figure 7. Differential gene expression heatmap for RP3 and R2.** The gene expression patterns of RP3 and R2 overlap as compared to the other clusters. In the top differentially expressed genes, R2 expresses 7 of the top 10 differentially expressed genes found in RP3.

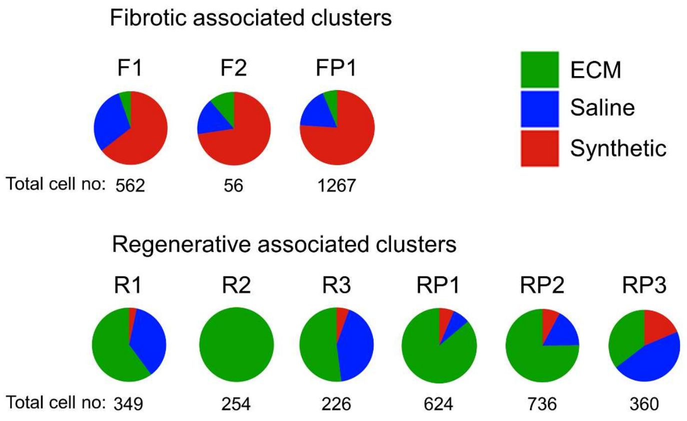


**Supplementary Figure 8. Cell composition of clusters by experimental condition.** The percent of cells per cluster is shown by experimental origin from regenerative, fibrotic, or saline conditions. Raw cell numbers from each condition were normalized to the total number of cells per condition.

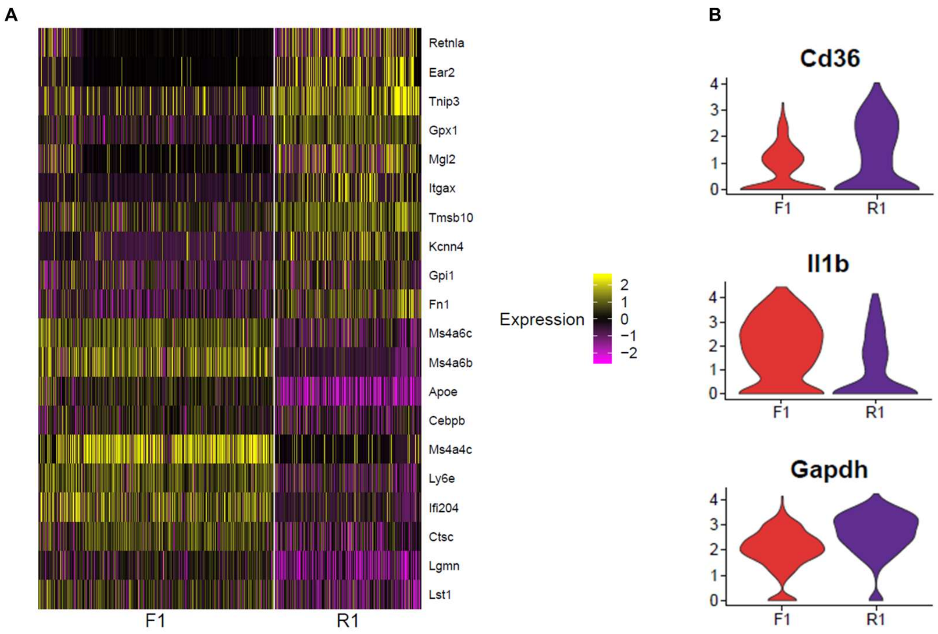


**Supplementary Figure 9. Expression of inflammatory genes in fibrotic cluster F2.** (A) Heatmap of top differentially expressed genes comparing F1 and R1. (B) Violin plots showing normalized expression of canonically inflammatory genes in F2.

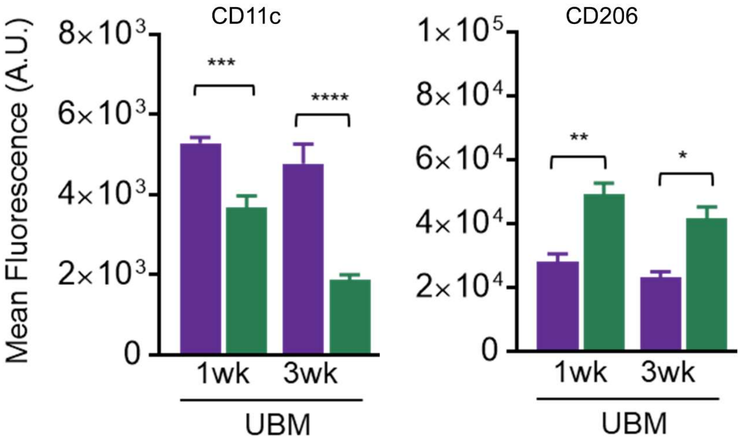


**Supplementary Figure 10. Quantification of expression levels of CD11c and CD206.** Protein expression of CD11c and CD206 for R1 and R2 at 1 and 3 weeks as quantified by flow cytometry.

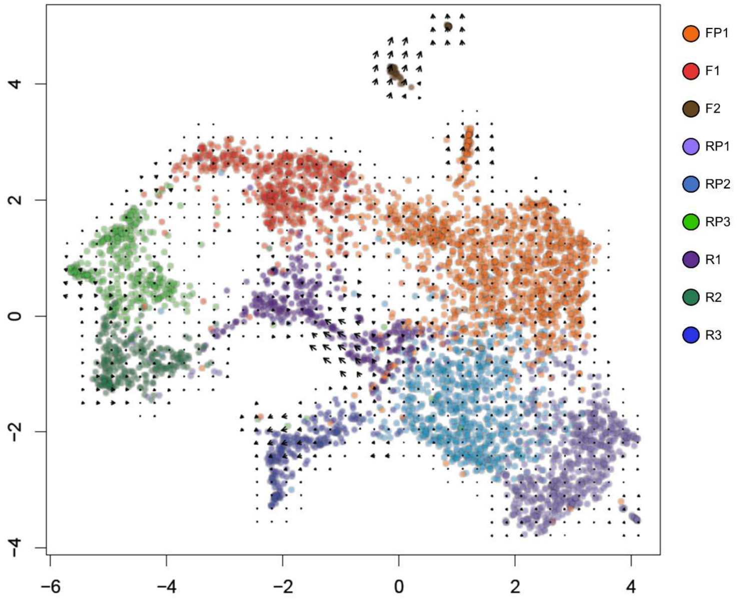


**Supplementary Figure 11. RNA velocity vector map.** RNA velocity vectors superimposed on a UMAP plot with cells colored by cluster.

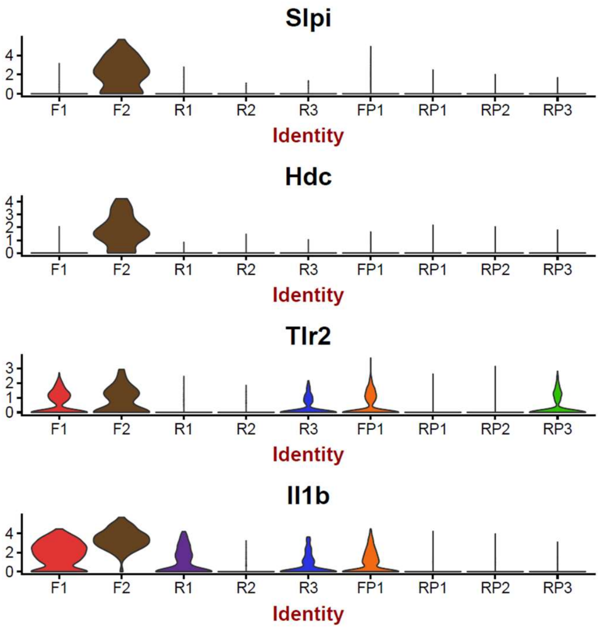


**Supplementary Figure 12. Expression of inflammatory genes in fibrotic cluster F2.** Violin plots show the normalized gene expression of canonical inflammatory genes in F2.

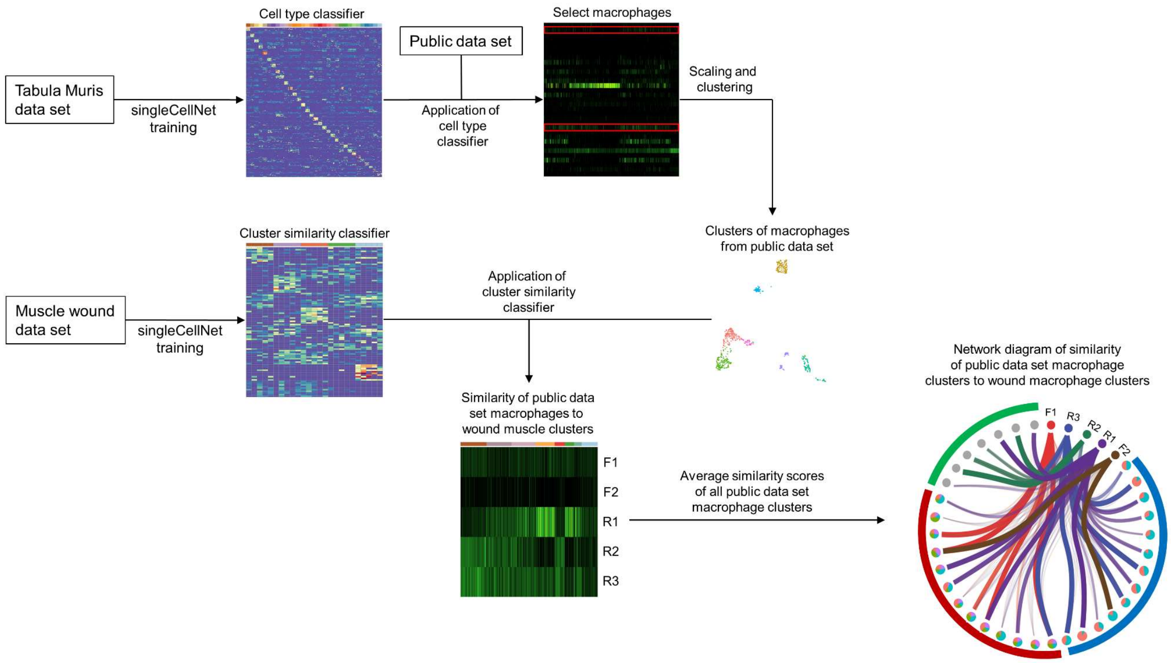


**Supplementary Figure 13. Analysis pipeline for publicly available data sets.** SingleCellNet was used to quantify similarity between macrophages derived from public data sets and macrophages from our wound microenvironment. First, SingleCellNet was used to generate a classifier for all cell types in the Tabula Muris data set, including macrophages and alveolar macrophages. The classifier was applied to each public data set, identifying cells with strong similarities to macrophages or alveolar macrophages. Cells classified in this manner were selected for further analysis and then scaled and clustered. In parallel, SingleCellNet was also used to generate a classifier from our five muscle wound terminal macrophage clusters. This classifier was then applied to the macrophages derived from the public data set, yielding five similarity scores for each cell in the public data set – one for each terminal macrophage cluster. These scores were visualized by heatmap. Finally, we averaged similarity score by each public data set cluster and used these values to generate a network diagram of similarities between terminal wound macrophage clusters and all clusters in the public data sets we analyzed.

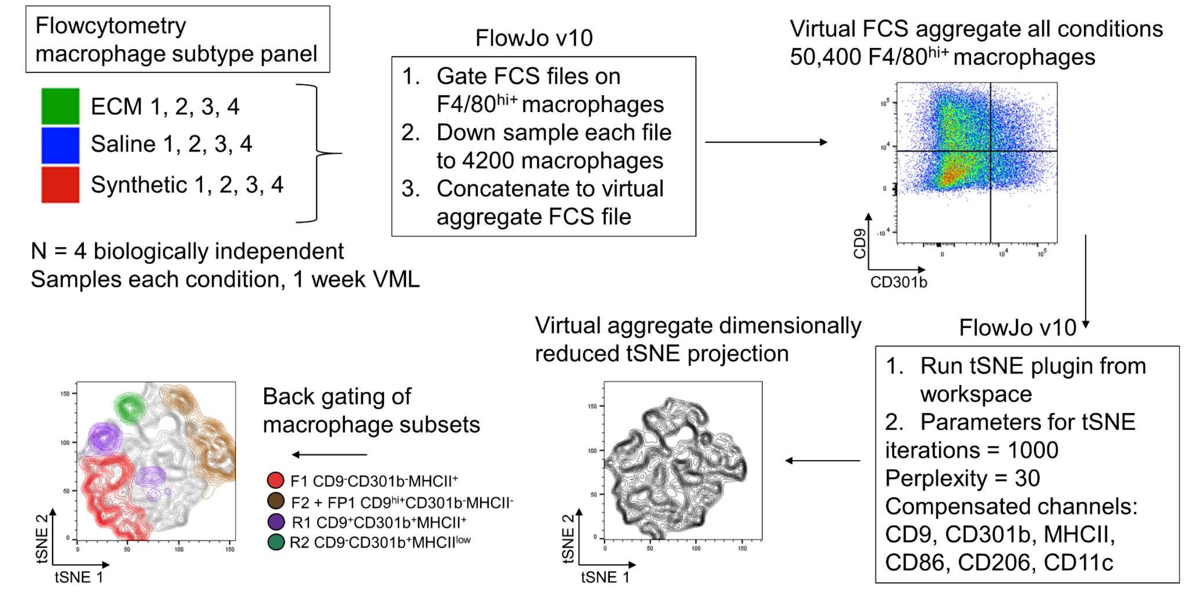


**Supplementary Figure 14. Scheme to generate for tSNE projection for virtual aggregate flow cytometry data.** Depicted is the multicolor flowcytometry method to generate a tSNE projection of fibrotic and regenerative macrophage subsets of *in vivo* samples.

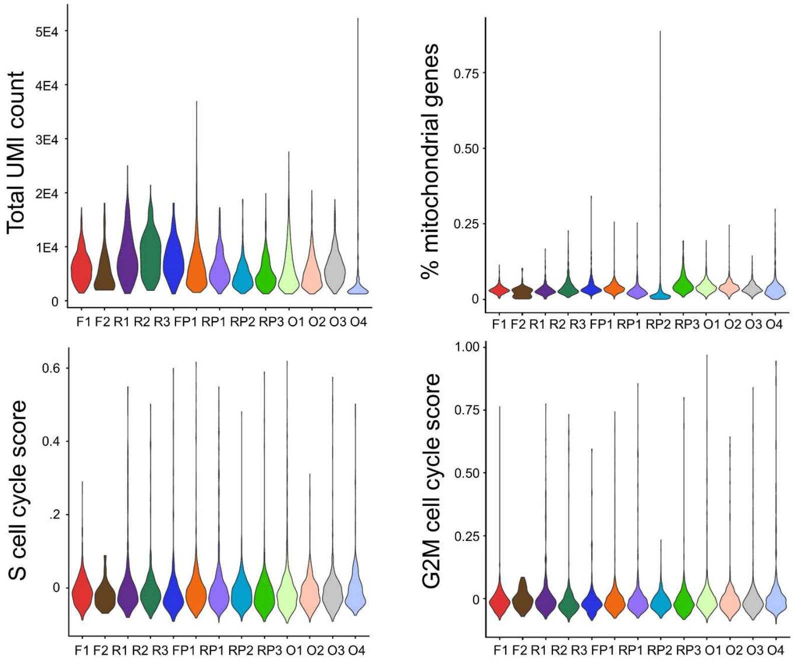


**Supplementary Figure 15. Scaling metric for the single cell RNA sequencing raw data.** Violin plots of all cell level metrics used in scaling including total UMI count, percent mitochondrial genes, Cell cycle S score, and cell cycle G2M scores.

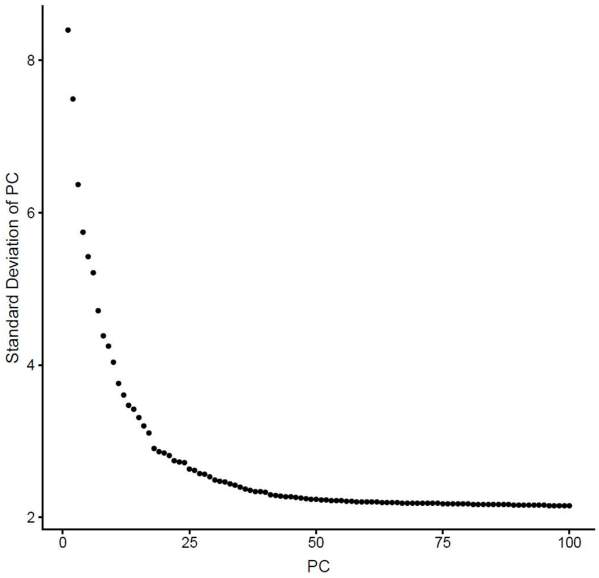


**Supplementary Figure 16. Variance by principle component.** Elbow plot shows the standard deviation of 50 principle components.

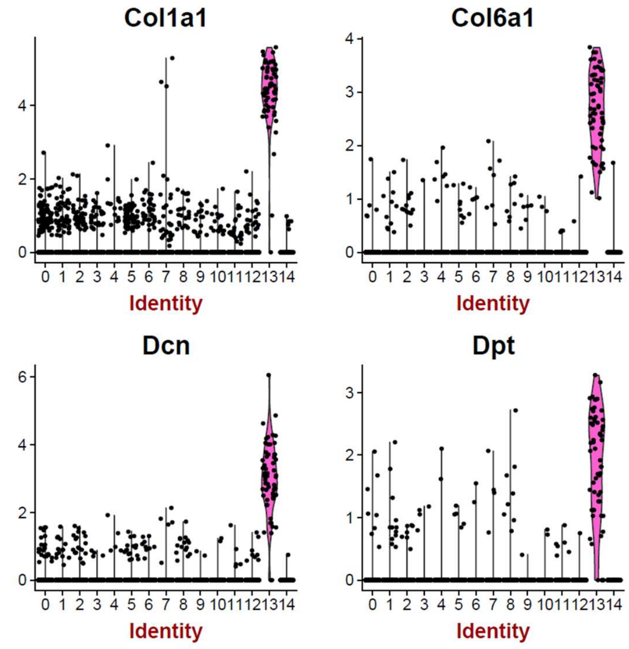


**Supplementary Figure 17. Variance by principle component.** Elbow plot shows the standard deviation of 50 principle components.

| | ECM | PCL | Saline |
|---|---|---|---|
| CellRanger estimated cell no. | 3394 | 2931 | 881 |
| Mean reads per cell | 91454 | 97594 | 56825 |
| Median genes per cell | 1741 | 1906 | 1528 |
| Fraction of reads in cells | 89.50% | 89.10% | 82.80% |
| Total genes detected | 15989 | 15648 | 13720 |
| Median UMI counts per cell | 5289 | 5438 | 3966 |
| Total read no. | 310M | 286M | 50M |
| Valid barcode % | 97.20% | 97.10% | 96.40% |
| Sequencing saturation | 89.60% | 88.70% | 83.60% |
| Q30 Bases in Barcode | 98.20% | 97.70% | 97.70% |
| Q30 Bases in RNA read | 92.50% | 92.70% | 92.60% |
| Q30 Bases in Sample index | 97.60% | 97.70% | 97.60% |
| Q30 Bases in UMI | 98.60% | 98.30% | 98.30% |

**Supplementary Table 1. Quality control statistics associated with alignment.** Quality control statistics from CellRanger alignment for each experimental condition.
